# Inducible Impairment of Polymerase Gamma Activity in Cardiomyocytes Promotes Severe Cardiomyopathy with Cardiac Hepatopathy

**DOI:** 10.64898/2026.02.17.706486

**Authors:** Simon T Bond, Yanie Tan, Shannen Walker, Sarah Jenkinson, Christine Yang, Yingying Liu, Kevin H Liu, Helen Kiriazis, Daniel G Donner, Jonathon Cross, Darren C Henstridge, David W Greening, Brian G Drew

**Author notes:** Authors for correspondence &.

## Abstract

Mitochondrial dysfunction is a hallmark feature of heart failure (HF) and cardiomyopathy, with substantial human and preclinical evidence suggesting that congenital mitochondrial defects can directly drive these conditions. Despite these strong links, not all individuals with mitochondrial disease develop cardiomyopathy, and therefore the precise mitochondrial defects that initiate cardiac pathology in this setting remain incompletely understood. Here, we describe a novel mouse model that induces progressive deterioration in mtDNA integrity specifically in cardiomyocytes. This model was generated through a post developmental, cardiomyocyte specific deletion of the exonuclease domain of Polymerase Gamma (PolG), impairing its ability to repair mtDNA. Strikingly, from just 16 weeks post-induction these mice displayed progressive worsening of cardiac output, ejection fraction, global strain and blood pressure culminating in premature death at 28-30 weeks post-mutation. Morphologically, these mice displayed cardinal features of hypertrophic cardiomyopathy with enlarged hearts, ventricles and cardiac fibrosis – but little evidence of congestive HF. We also demonstrate using various transcriptional and proteomic readouts, that signalling pathways characteristic of cardiomyopathy were activated. Interestingly, prior to substantial declines in cardiac function, we detect robust activation of the integrated stress response (ISR) in PolG^Mut^ mice, and major rewiring of mitochondrial folate metabolism pathways, suggesting that these pathways underpin the developing pathology. Lastly, we also observed a striking hepatopathy phenotype in mutant mice reminiscent of that observed in patients with HF, a condition not robustly recapitulated previously in other mouse models. Thus, our data establish a direct link between mtDNA instability leading to chronic activation of the ISR and rewiring of folate metabolism in cardiomyocytes, subsequently driving a severe cardiac phenotype associated with hepatopathy.

## INTRODUCTION

Mitochondria are well known for their role in generating ATP, however, mounting evidence highlights additional complex and diverse roles for mitochondria in a variety of cellular processes and pathologies. One of the unique characteristics of mitochondria is that they are the only mammalian organelle other than the nucleus, to harbour their own genome. This mitochondrial DNA (mtDNA) is essential for the effective functioning of oxidative phosphorylation machinery and is particularly vulnerable to damage due to its limited protection from the mitochondrial matrix and proximity to reactive oxygen species. Fortunately, several pathways have evolved in mammalian cells to protect mtDNA from these deleterious effects, which when working efficiently can effectively preserve mtDNA integrity – however when they fail, disease can ensue.

Mitochondria are especially critical in cardiomyocytes, due to the heart’s high-energy demand and thus reliance on mitochondrial function to maintain contractility and overall function. Disruptions to mitochondrial processes, including defects in mtDNA replication, protein import, reductions in mitochondrial abundance or altered mitochondrial dynamics, have been linked to the pathogenesis of cardiometabolic diseases^1–5^. Reductions in mitochondrial activity that result in energy deficits, can have a significant effect on cardiac function and output, however, are not the only causal drivers. One key mitochondrial signalling axis that has recently attracted significant interest, is the mitochondrial integrated stress response (mtISR)^6,7^. This adaptive response is triggered by various forms of mitochondrial dysfunction and orchestrates transcriptional programs aimed at acutely restoring cellular homeostasis^7,8^. Central to this response is the activation of the kinase eIF2α, which is activated by phosphorylation at serine 51 (Ser51), promoting a cascade of events that ultimately leads to the induction of nuclear-encoded genes that support metabolic rewiring, amino acid metabolism, and redox balance^9,10^. Moreover, mtISR activation, particularly in muscle tissues, leads to the systemic release of mitokines such as growth differentiation factor 15 (GDF15) and fibroblast growth factor 21 (FGF21), which are thought to act as endocrine signals in response the ISR to modulate metabolism or liberate energy stored elsewhere in peripheral tissues^11^.

Despite growing interest in the mtISR, its activation and the subsequent functional changes that occur in the heart in response to this pathway, remain incompletely understood. In particular, it is not known what specific precipitating events lead to ISR activation in the heart, and whether it is chronic activation of the ISR that ultimately promotes mitochondrial dysfunction and disease, or vice versa. To address this gap, we generated a conditional Polymerase Gamma (PolG) in-frame truncation^11^ model in which mtDNA integrity is disrupted selectively in cardiomyocytes of affected mice (PolG^Mut^). PolG is the sole polymerase found in the mitochondria, where it is responsible for replicating and repairing mtDNA. Accordingly, inefficiencies in PolG activity lead to a vast array of degenerative and often fatal conditions in humans, whilst preclinical models of PolG disease present with similar neuro-, musculo- and cardio-degenerative phenotypes^11–17^.

Herein, we demonstrate that cardiomyocyte specific mtDNA damage induced by PolG truncation, leads to robust and early activation of the mtISR in the heart, leading to progressive hypertrophic cardiomyopathy and rapid deterioration of cardiac function that ultimately culminates in premature death. This phenotype appears to be preceded by major rewiring of 1-carbon metabolic processes in cardiomyocytes, specifically folate metabolism in the mitochondria, pointing to a potential pathway of interest for subsequent therapeutic intervention. Moreover, this model also developed the unique complication of cardiac hepatopathy, which is observed in a subset of patients living with decompensated cardiomyopathy, but has not been readily recapitulated in preclinical models. Thus, this model represents a unique platform to dissect the molecular mechanisms linking mtDNA instability, mtISR signalling, and cardiac pathology, revealing potential novel therapeutic targets for the treatment of ISR driven cardiomyopathies.

## RESULTS

### Cardiomyocyte specific PolG impairment leads to mtDNA damage, acute weight loss, cardiac hypertrophy and premature death

In previous published work from our group, we have demonstrated that inducible, skeletal muscle specific truncation of the PolG exonuclease domain (in-frame loss of exons 4 & 5), leads to progressive muscle wasting, loss of body fat and ultimately a cachexia like phenotype over a 12 month period^11^. This was associated with a chronic activation of the ISR pathway, and subsequent rewiring of metabolic pathways that preceded disease development^11^. Accordingly, in the current study we sought to determine the impact of this same truncation in cardiomyocytes, and determine the effects of cardiac specific mtDNA mutations/deletions on heart function and whole-body health. Whilst previous studies have investigated the impact of developmental single point mutants of PolG in the heart^18^, our model is the first to investigate PolG impairment in an inducible, tissue specific, post-developmental manner – mimicking an “acquired” like disease phenotype in the heart, as opposed to a congenital condition.

To do this we crossed our novel PolG^fl/fl^ mice^11^ with MHCα Cre-ERT2 mice to generate an inducible cardiomyocyte specific PolG exonuclease mutant mouse, that would allow for a post-developmental increase in mtDNA mutations/deletions explicitly in cardiomyocytes (**Figure 1A**). To initially understand the broader impacts on health in this model, we designed a study where at 8 weeks of age (WOA) half of the mice in cohorts of fl/fl and fl/fl+cre mice were given vehicle (oil, PolG^Con^), and the other half were given tamoxifen (oil+TAM) to induce PolG mutation. These mice were then proposed to be characterised for up to 12 months. However, a severe phenotype developed by ∼30 weeks in the fl/fl+cre mice that received tamoxifen (PolG^Mut^), including rapid weight loss from 25 weeks post-TAM (**Figure 1B**, **Supplemental Figure 1A**), culminating in premature death in these animals at ∼28-30 weeks post-TAM (**Figure 1C**). Indeed, the mutant animals appeared smaller in size when observed at the whole-body level (**Figure 1D**). The weight loss observed in mutant animals was mostly fat mass (**Supplemental Figure 1B&C**), with only minor changes in lean mass (**Supplemental Figure 1D&E**). There was also a significant change in organ weights at study end in epididymal fat (Epi), subcutaneous fat (SubQ), brown adipose tissue (BAT) and liver in PolG^Mut^ mice, but no change in quadricep muscle (Quad) (**Supplemental Figure 1F**).

**Figure 1:**
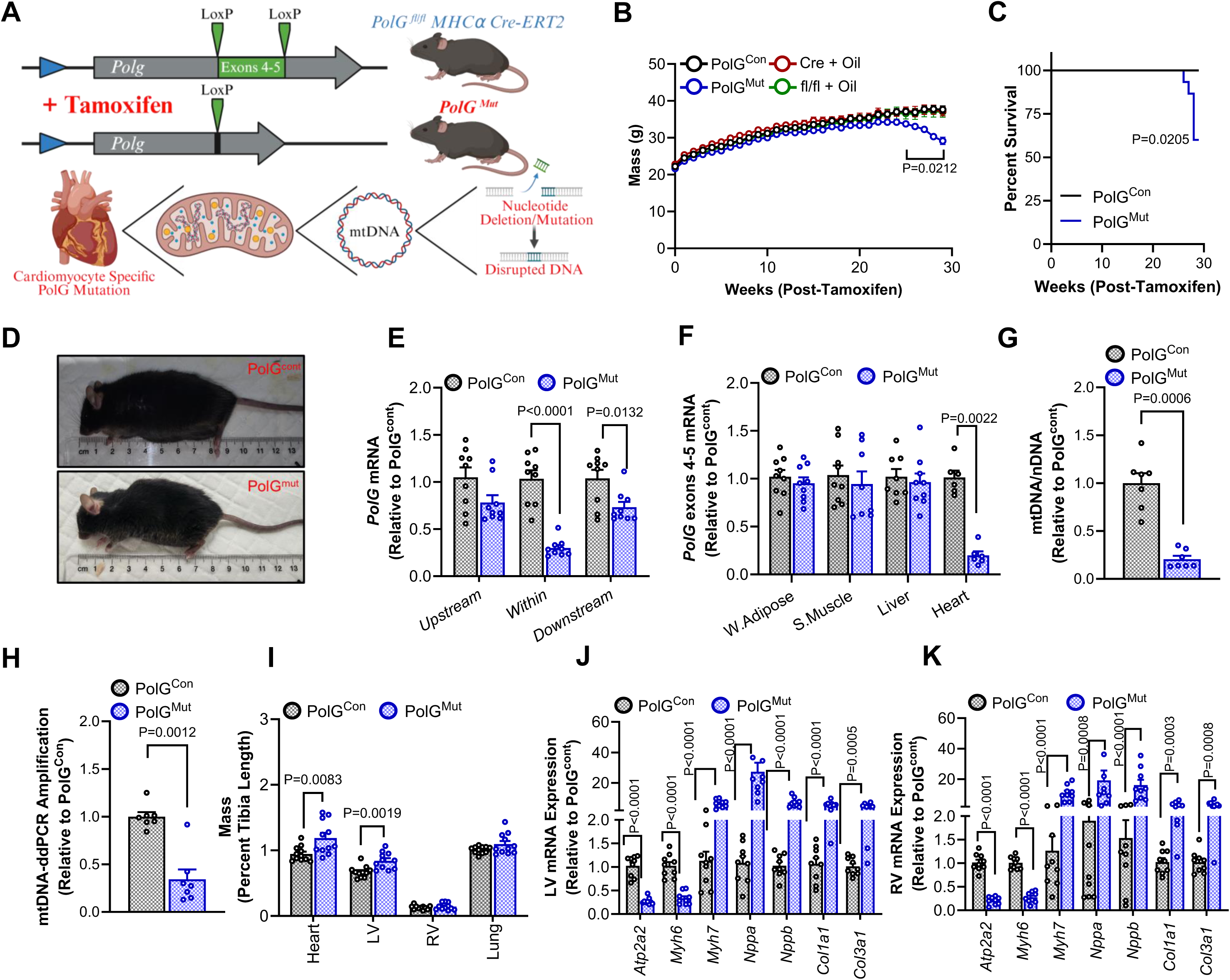
Cardiomyocyte specific PolG mutation leads to acute weight loss and premature death. **(A)** Schematic of tamoxifen induced cardiomyocyte specific excision of *Polg1* exons 4-5 and subsequent mtDNA damage. **(B)** Weekly whole-body weights in PolG^Mut^ (PolG^fl/fl^ MHCα Cre-ERT2 + tamoxifen, n=15), PolG^Con^ mice (PolG^fl/fl^ MHCα + tamoxifen, n=11), Cre+Oil (MHCα Cre-ERT2 + oil, n=6) and fl/fl+Oil (PolG^fl/fl^ MHCα Cre-ERT2 + oil, n=5) mice. **(C)** Kaplan-Meier graph indicating percentile survival rate in PolG^Mut^ (n=15) and PolG^Con^ (n=11) mice with **(D)** representative images at 30 weeks post-Tam. **(E)** Left ventricle (LV) PolG gene expression measured by qPCR at *Polg1* regions pre-exon 4, within exons 4-5 and post-exon 5 (n=9/group). **(F)** PolG exons 4-5 qPCR gene expression in white epididymal adipose (W.Adipose, n=9/group), quadricep skeletal muscle (S.Muscle, n=9/group), liver (PolG^Mut^ n=8 and PolG^Con^ n=9) and heart (n=6/group) from PolG^Mut^ and PolG^Con^ mice. **(G)** Mitochondrial DNA (mtDNA) to Nuclear DNA (nDNA) ratio (relative to control, n=7/group) and **(H)** amplification of mtDNA region (relative to control, n=7/group) in PolG^Mut^ and PolG^Con^ mice LV at 30 weeks post-Tam. **(I)** Wet tissue weights normalised to tibia length at 30 weeks post-Tam (n=11/group). Expression of genes associated with heart disease in **(J)** LV and **(K)** RV (right ventricle) from PolG^Mut^ and PolG^Con^ mice (n=9/group). Data are presented as mean ± SEM, with P-value between control and mutant biological replicates determined by repeated measures two-way ANOVA with correction for multiple comparisons **(B)**, Mantel-Cox test **(C)**, unpaired welch t-test **(E)** or Mann-Whitney test **(F-K)**. Source data for these figures are provided in the Source Data file. Fig 1A created in BioRender. Drew, B. (2025), http…….

Subsequent molecular analysis of tissues from PolG^Mut^ and PolG^Con^ mice confirmed that our model demonstrated a significant reduction in exons 4-5 of the PolG mRNA in left ventricular (LV) tissue (**Figure 1E**: “within” the exon 4-5 floxed region, compared with regions “upstream” and “downstream” of exons 4-5), and that this was specific to cardiac tissue, demonstrated by reductions in *Polg* mRNA expression in heart but not adipose, liver or quadricep (**Figure 1F**). Although we demonstrate modest but significant reductions in overall cardiac *Polg* mRNA expression, this is comparable to global PolG mutator models and our previous muscle specific model^11^, and importantly does not suggest a complete “deletion” of the mRNA. As a consequence of this mutation in PolG, we show that our model has impacts on mtDNA abundance in heart tissue, through a reduction in the mitochondrial DNA to nuclear DNA (mtDNA/nDNA) ratio (**Figure 1G**), and also a reduced amplification of the D-loop region from precise amounts (1ng) of mtDNA between PolG^Mut^ and PolG^Con^ hearts, suggesting an increase in mtDNA deletions (and thus loss of amplification signal) in mutant heart tissue (**Figure 1H**).

To investigate the pathology and molecular underpinnings of disease in the hearts of our mutant model, we dissected tissues at end point (30 weeks post-TAM) and analysed heart and lung weights to investigate phenotypic aspects of cardiomyopathy and heart failure. Indeed, we demonstrate that heart weight and left ventricle (LV) weight in these animals was increased compared to PolG^Con^ mice, but right ventricle (RV) and lung weights were not changed (**Figure 1I**), indicating a predominantly left heart related cardiomyopathy that was likely not congestive. When we investigated molecular pathways associated with cardiomyopathy using qPCR in both LV (**Figure 1J**) and RV (**Figure 1K**) tissue, we observed striking alterations in all of these genes in both tissues of mutant mice, confirming a robust cardiomyopathy phenotype in this model.

With the observation of a fatal cardiomyopathy observed in our first cohort of PolG^Mut^ mice by approximately 30 weeks post-TAM, we next wanted to understand the functional and temporal nature of this phenotype, and therefore we generated several additional cohorts to investigate the model at varying timepoints along the disease progression timeline (**Figure 2A**). These studies were designed to acquire cardiac functional data across 4, 16, 20, 24 and 28 weeks post-TAM, as well as tissue collections at 20, 24 and 28 weeks, which are time points that reflect early, mid and late cardiomyopathy presentation.

**Figure 2:**
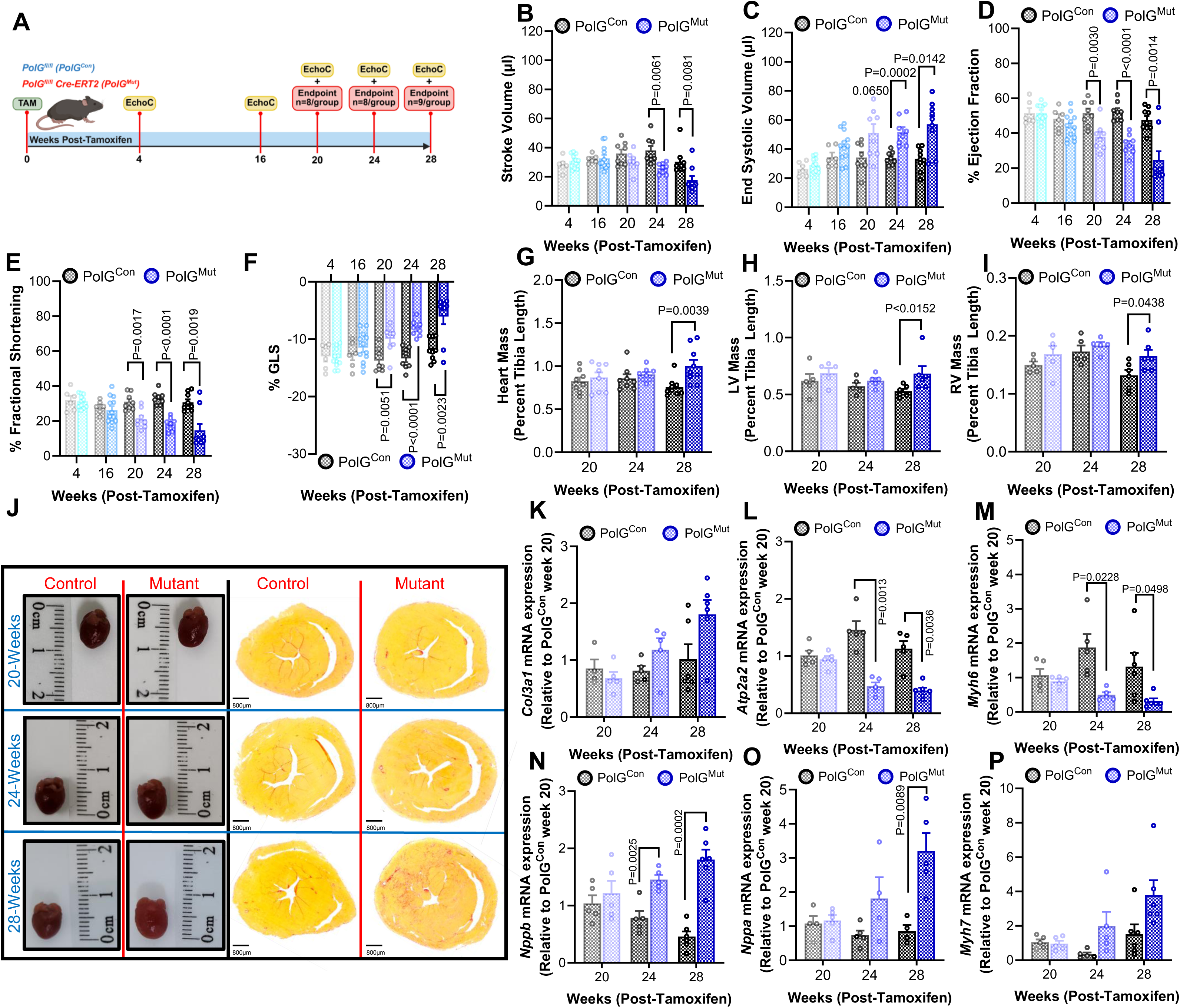
Mitochondrial DNA damage in cardiomyocytes promotes cardiac remodelling and impacts heart function. **(A)** Schematic outlining animal study to investigate heart function from 0-28 weeks post-Tam. Cardiac function, including **(B)** stroke volume, **(C)** end systolic volume, **(D)** percent ejection fraction, **(E)** percent fractional shortening and **(F)** percent GLS (global longitudinal strain) was assessed by echocardiography at 4 (PolG^Mut^ n=11 and PolG^Con^ n=6), 16 (PolG^Mut^ n=12 and PolG^Con^ n=6), 20 (n=8/group), 24 (n=8/group) and 28 (n=9/group) weeks post-Tam in PolG^Mut^ and PolG^Con^ mice. Wet tissue weight normalised to tibia length in **(G)** Heart, **(H)** LV and **(I)** RV from PolG^Mut^ and PolG^Con^ mice (20, 24 week n=8/group and 28 week n=9/group). **(J)** Representative images of PolG^Mut^ (Mutant) and PolG^Con^ (Control) mice whole hearts and heart sections stained with picrosirius red at 20, 24 and 28 weeks post-Tam. Expression of genes associated with **(K)** fibrosis and **(L-P)** heart disease (relative to PolG^Con^ mice at 20 weeks post-Tam), as determined by qPCR in LV from PolG^Mut^ and PolG^Con^ mice (20, 24 week n=5/group and 28 week n=6/group). Data are presented as mean ± SEM, with P-value between control and mutant biological replicates determined by unpaired welch t-test **(B, D-I, K-P)** or Mann-Whitney test **(C)**. Source data for these figures are provided in the Source Data file. Fig 2A created in BioRender. Drew, B. (2025), http…….

Echocardiography data demonstrate that there was a significant and consistent impact on cardiac function from 20 weeks post-TAM, although there was a trend for a difference in some functional readouts from as early as 16 weeks post-TAM. This can be observed in parameters including Stroke Volume (**Figure 2B**), End Systolic Volume (**Figure 2C**), Ejection Fraction (**Figure 2D**), Fractional Shortening (**Figure 2E**) and Global Longitudinal Strain (GLS) (**Figure 2F**). This is mostly consistent with other echocardiography read outs including Cardiac Output (**Supplemental Figure 2A**) and End Diastolic Volume (**Supplemental Figure 2B**). Furthermore, when analysing tissue weights in mice at end points of 20, 24 and 28 weeks post-TAM, we observed a significant increase in heart mass (**Figure 2G**), LV mass (**Figure 2H**) and RV mass (**Figure 2I**) only at the 28 weeks post-TAM timepoint, although there were trends for an increase at all time points. These data are consistent with echocardiography estimated Left Ventricular Mass (LVM) at both end diastole (EDLVM; **Supplemental Figure 2C**) and end systole (ESLVM; **Supplemental Figure 2D**). These data provide strong evidence for an underlying cardiomyopathy phenotype that impacts on heart function, but that decompensation of the myopathy only occurs in the late stage of the disease (i.e. 28 weeks post-TAM). Consistent with hemodynamic changes and functional effects of the mutant earlier in the phenotype, we observed that heart rate was elevated at several time points, including the earlier time point of 16 weeks (**Supplemental Figure 2E**), with Systolic (**Supplemental Figure 2F**) and Diastolic (**Supplemental Figure 2G**) blood pressure also demonstrating differences at some of the earlier time points. Such changes are consistent with HR increasing in an attempt to maintain cardiac output, as stroke volume was likely decreasing due to mild defects in heart function.

With the obvious deterioration in cardiac function in the mutant mice, we were interested in observing if there were any changes in structure and fibrosis in the heart. Accordingly, we analysed sections of hearts from 20, 24 and 28 week control and mutant mice, and demonstrated that consistent with tissue weight and echo data, heart size was increased at 28 weeks (**Figure 2J**), and also demonstrated a significant increase in fibrosis of the LV and RV (**Figure 2J**) and observed a trend for increased expression of the collagen related gene *pro-collagen* 3 in the mutant hearts over time (**Figure 2K**). In line with the presentation of cardiac hypertrophy and dysfunction, expression of genes characteristic of the development of cardiomyopathy were also altered in the mutant hearts across the time course. Specifically, we observed a significant change in *Atp2a2* (SERCA2) (**Figure 2L**), *Myh6* (**Figure 2M**) and *Nppb* (BNP) (**Figure 2N**) at 24 and 28 weeks, *Nppa* (ANP) (**Figure 2O**) at 28 weeks, and a trend for increased *Myh7* (**Figure 2P**) at 28 weeks. These gene expression patterns are consistent with a maladaptive cardiac pathology and are commonly observed in models of hypertrophy and cardiomyopathy.

Collectively, these results suggest that PolG^Mut^ mice maintain a healthy phenotype, with regular heart function up to ∼20-weeks post-TAM, upon which an accumulation of mtDNA damage leads to rapid cardiac remodelling and weight loss that results in impaired cardiac function and premature death.

### Molecular Pathways Driving Severe Heart Failure in PolG Mutant Hearts

Having confirmed in the original 30 week study that mtDNA damage was indeed apparent in our mutant mice (Figures 1G&H), we next sought to understand what cardiomyocyte specific pathways were being impacted by this, both early and late, that might be responsible for inducing such rapid and striking phenotypes. The obvious candidate for such direct effects on the heart is impaired mitochondrial activity driven by accumulating mtDNA damage. However, before analysing mitochondrial functional readouts, it was important to demonstrate the spectrum and degree of mtDNA damage over time, as this could provide relevant insights into the potential downstream molecular pathways that might be impacted in our model.

We therefore sequenced the mtDNA from hearts of mice at the 20 and 28 week post-TAM time points (prior to and after changes in heart function). These data clearly demonstrate that our PolG exonuclease mutant develops numerous and varied mutations in the mtDNA, even at 20 weeks, and that these mutations continue to accumulate as the model progresses toward end-stage. This can be observed in the mtDNA mutation frequency plot (**Figure 3A**), demonstrating that mtDNA from mutant mice have a higher frequency of mutations in specific regions of the sequence (namely between 1-8000bp of the mitochondrial genome), which increase further in the 28-week animals. In terms of the type of mutation occurring, the substitution spectrum plot (**Figure 3B**) demonstrates that mutant mice develop a progressively increasing mix of point mutations and insertions/deletions (indels), indicating an expected heterogenic mutation pattern, due to an impaired PolG activity in cardiomyocytes of PolG^Mut^ mice. This outcome is important, as it has previously been shown that high point mutation burden in mtDNA leads to defects in OXPHOS activity^13,19^, whereas indels often lead to activation of stress responses such as the integrated stress response (ISR)^9,20^.

**Figure 3:**
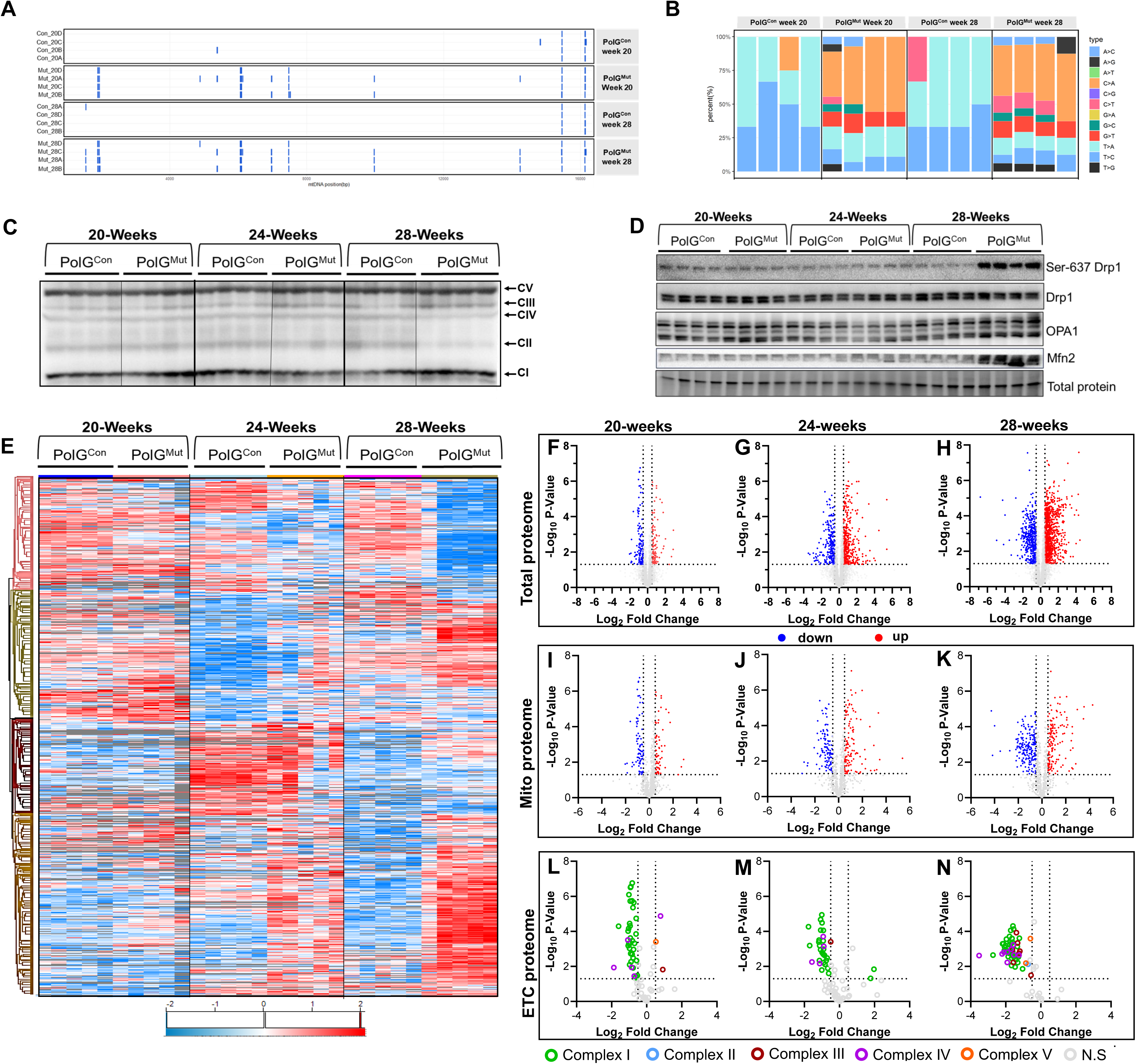
Progressive regulation of the cardiac proteome induced by mitochondrial DNA mutations/deletions. **(A)** Mitochondrial mutation load and mutation base pair location (blue lines indicating changes from the NCBI C57BL/6J mtDNA reference sequence) at 20 and 28 weeks post-Tam in mtDNA of LV from PolG^Mut^ and PolG^Con^ mice (n=4/group). **(B)** Ratio of mutation type in PolG^Mut^ compared with PolG^Con^ mice (n=4/group). **(C)** Immunoblot for mitochondrial OXPHOS proteins from CI, CII, CIII, CIV and CV and **(D)** mitochondrial dynamics regulatory proteins including phosphorylated Drp1 (Ser-637 pDrp1), total Drp1 (Drp1), Optic Atrophy mitochondrial dynamin like GTPase 1 (OPA1) and Mitofusin 2 (Mfn2) in LV from PolG^Mut^ and PolG^Con^ mice at 20, 24 and 28 weeks post-Tam (n=4/group/timepoint). Unbiased assessment of protein abundance analysed from left ventricle (LV) of PolG^Mut^ and PolG^Con^ mice at 20, 24 and 28 weeks post-Tam (n=5/group/timepoint) and computational analysis of the data was performed including **(E)** hierarchal clustering of differentially expressed proteins and volcano plots of significantly differentially expressed proteins (grey=not significant) for **(F-H)** total proteome, **(I-K)** mitochondrial proteins and **(L-N)** electron transport chain (ETC) proteins. Protein intensities were log2 transformed and normalized using quantile normalization from R package preprocess Core and proteins were subjected to PCA and student’s t-test. Source data for these figures are provided in the Source Data file.

Knowing that there were substantial point mutations present in the mtDNA of our mutant mice, we were interested to know if this was impacting on mitochondrial function at different time points. Therefore, given the major role that OXPHOS plays in regulating respiration and ATP production in the mitochondria, we first performed Western blotting for abundance of ETC complexes in the LVs of control and mutant mice (**Figure 3C**). These data demonstrate that there was no difference observed in these readouts until 28-weeks post-TAM, when there are significant reductions in Complex II and Complex IV, with no detectable change in any other Complexes. With these late changes observed, it was suggestive to us that this may be related to the HF phenotype itself, as opposed to being the causal driver of the pathology. Therefore, we next performed additional Western blots to investigate the status of mitochondrial fission/fusion pathways (**Figure 3D**), as these have often been shown to be sensitive to subtle changes in mitochondrial function^21,22^. These data also demonstrate that there was minimal change in these pathways at 20- and 24-weeks, but significant change at 28-week post-TAM mice. Collectively, these data suggest that significant mitochondrial dysfunction does not develop until later in the phenotype (28 weeks), and is likely to be secondary to other more primary molecular changes. To investigate this more broadly, we performed proteomic analysis on LVs from mice at 20, 24 and 28-weeks post-TAM, to gain a more global understanding of the molecular changes occurring in mutant hearts.

Proteomics was performed on multiple samples (n=5) from each group and time point, demonstrating a robust and consistent detection of >3000 proteins in all groups (**Supplemental Figure 3A-B**). Principle component analysis confirmed a high level of clustering of the individual group samples (**Supplemental Figure 3C**). When analysing the data for differential abundance, we observed significant and varied changes in the proteome compared to control mice across all three time points studied (20, 24 and 28 weeks post-TAM). This can be readily observed in the heatmap and hierarchical clustering (**Figure 3E**), which highlights that there were specific molecular pathways being altered at each time point, providing interesting insights into those which might be causal to the phenotype. To investigate this in more detail, we performed a pathway enrichment analysis on the protein-sets increased and decreased (compared to control mice) at each time point (**Supplemental Figures 3D-I**). Two interesting observations arose from the pathway enrichment analyses. Firstly, there was broad identification of several disease phenotypes (i.e. Parkinsons, Huntingtons, Alzheimers, fatty liver) being enriched in the downregulated pathways at 20-weeks, including oxidative phosphorylation (i.e. ETC abundance). This is consistent with mild mitochondrial alterations, which remained in the later time points but was equally matched with pathways consistent with cardiomyopathy. Secondly, was the enrichment of stress pathways and alterative metabolism in the upregulated networks at all time points, particularly those at the 20-week time point. These pathways include AA metabolism, tRNA biosynthesis and carbon metabolism, all suggesting that there were early-stage stress pathways being activated, and metabolic rewiring occurring in these early stages.

To investigate the molecular phenotypes, including mitochondrial networks, in more detail, we plotted out the overall changes in proteome that were occurring at the different time points. This can be seen firstly for total proteome (**Figure 3F-H**), which shows progressively increasing differential expression as the HF phenotype progresses. When we specifically observe the mitochondrial proteome (**Figure 3I-K**), we see a similar pattern as the total proteome, with only a few regulated proteins at 20 weeks, but a substantial number at 28-weeks. These data are consistent with those shown above in Western blots, which demonstrate that major changes in mitochondrial proteins only begin to become apparent in the later stages of pathology. Finally, we plot the ETC proteome (Complexes I-V) (**Figure 3L-N**), and demonstrate that for the most part there are only mild (less than 2-fold) reductions in Complex I proteins apparent at 20 weeks, but substantial (upto 4-fold) reductions in many proteins associated with Complex I, II and IV apparent at 28-weeks. Collectively, these data support the notion that in the early stages of pathogenesis in this mutant model, there is activation of stress and alternative pathways, and the beginnings of mitochondrial dysfunction, all of which precede molecular and functional features of cardiomyopathy. Considering these findings, we were interested in identifying what stress and metabolic pathways may be changed in these early time points, in order to determine if these might be precipitating the cardiomyopathy phenotype.

From previous work in our lab, and others, it has been shown that mtDNA mutations can activate a pathway known as the integrated stress response (ISR). Indeed, prior work from our lab has shown in a skeletal muscle specific model of PolG exonuclease mutation, that the ISR is an early and chronic pathway activated in this tissue^11^. Proteomic data shown from hearts in the current study, demonstrates that early pathways being changed include AA metabolism, tRNA biosynthesis and carbon metabolism – all of which are characteristic of ISR activation. Hence, we next set out to determine if an early and precipitating event in the HF phenotype of these mice, might be related to chronic activation of the ISR.

The critical early molecular event in the propagation of ISR activity is phosphorylation of the kinase eIF2α at serine 51 (Ser51 p-eIF2α). We therefore performed Western blots for phosphorylation of eIF2α in cardiac tissue (LV) as a measure of ISR activation in our mouse model. We first detected significant increases in both phosphorylated and total eIF2α in hearts from the most end-stage mice from our studies, being 30-weeks post-TAM (**Figure 4A-B)**. These data importantly implicate ISR activation in failing hearts from our mutant model, but cannot determine if this activation is related to PolG induced mtDNA mutations, or if it is just part of the overall end-stage HF response. Accordingly, we further performed Western blots in the hearts from mice at 20-, 24- and 28-weeks post-TAM (**Figure 4C-D)**. These data demonstrate that there is already significant phosphorylation of eIF2α at the 20-week time point, which becomes progressively more prominent at 24 and 28-weeks post-TAM.

**Figure 4:**
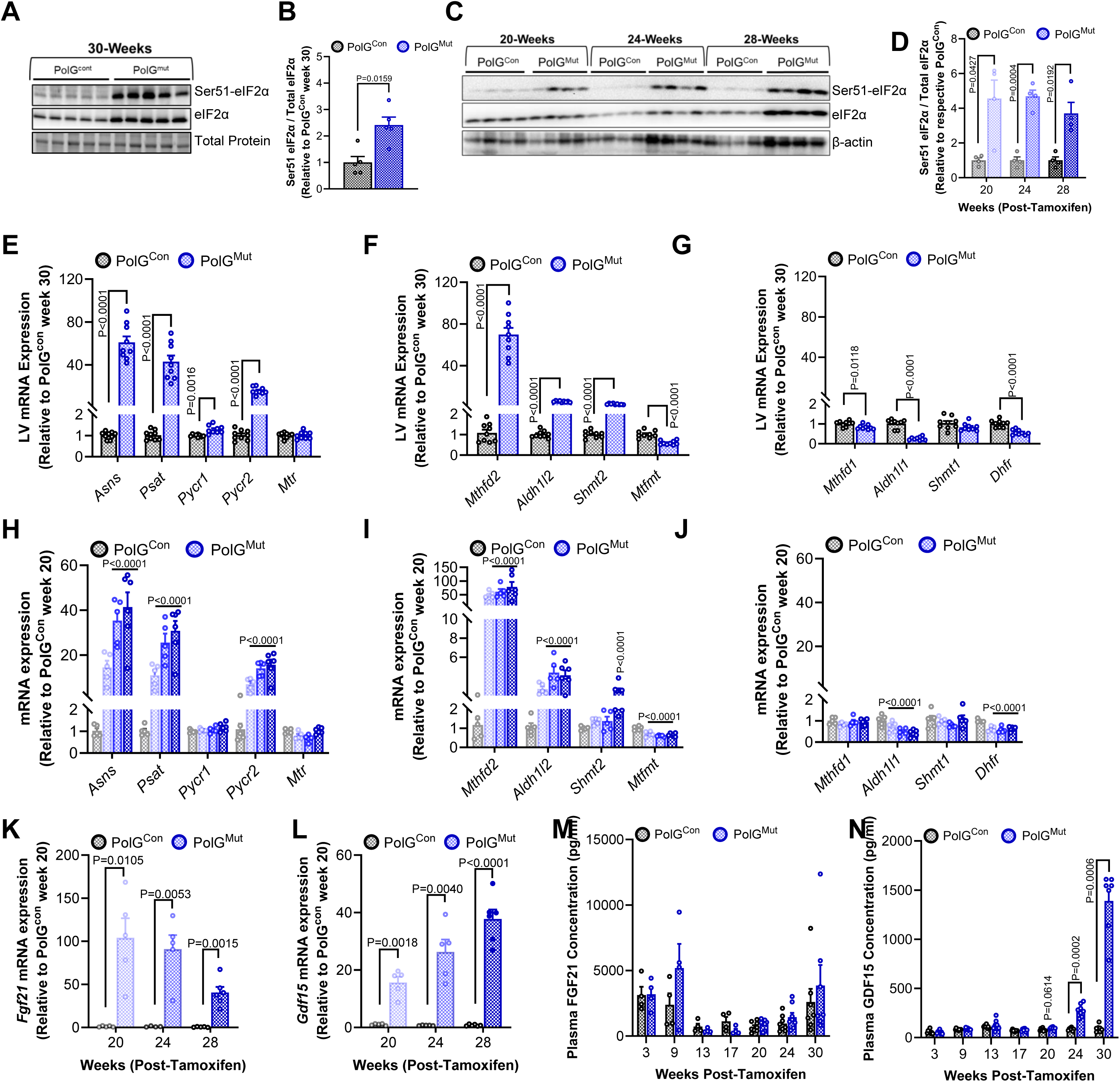
Accumulating mitochondrial DNA damage in the heart activates the integrated stress response. **(A)** Immunoblot of phosphorylated and total eIF2α in LV from PolG^Mut^ and PolG^Con^ mice at 30 weeks post-Tam (n=5/group), with **(B)** corresponding densitometry ratio of phosphorylated to total eIF2α. **(C)** Immunoblot of phosphorylated and total eIF2α was repeated in LV from PolG^Mut^ and PolG^Con^ mice at 20, 24 and 28 weeks post-Tam (n=4/group/timepoint) with **(D)** corresponding densitometry ratios. Expression of genes (qPCR) associated with **(E)** amino acid and 1-Carbon metabolism, and genes associated with folate metabolism within **(F)** the mitochondria and **(G)** the cytosol, in LV from PolG^Mut^ and PolG^Con^ mice at 30 weeks post-Tam, (n=9/group). Gene expression was repeated in LV from PolG^Mut^ mice at 20 (n=5), 24 (n=5), and 28 weeks post-Tam (n=6) relative to PolG^Con^ mice at 20 weeks post-Tam (n=5) for **(H)** amino acid and 1-Carbon metabolism, and genes associated with folate metabolism within **(I)** the mitochondria and **(J)** the cytosol. **(K)** *Fgf21* and **(L)** *Gdf15* gene expression (qPCR) in LV from PolG^Mut^ mice at 20 (n=5), 24 (n=5) and 28 (n=6) weeks post-Tam and PolG^Con^ mice at 20 (n=5), 24 (n=5) and 28 (n=5) weeks post-Tam (relative to PolG^Con^ mice at 20 weeks post-Tam). Plasma protein concentration of **(M)** FGF21 and **(N)** GDF15 in PolG^Mut^ and PolG^Con^ mice at 3, 9, 13, 17, 20, 24, and 30 weeks post-Tam (FGF21: 3, 9, 13, 17 weeks n=4, 20, 24 weeks n=8, 30 weeks n=7; GDF15: 3, 9, 13, 17, 20, 24 weeks n=8, 30 weeks n=7). Data are presented as mean ± SEM, with P-value between control and mutant biological replicates determined by two-way ANOVA with correction for multiple comparisons **(H-J: P-value compared with PolG^Con^ mice at 20 weeks post-Tam)**, unpaired welch t-test **(D, E, G, K, L)** or Mann-Whitney test **(B, F, M, N)**. Source data for these figures are provided in the Source Data file.

To further investigate the level of ISR activation at these different time points, we used qPCR data to investigate several genes that are known to be transcriptionally regulated by the ISR, which include genes from the 1-Carbon metabolism pathway (1C-Met), amino acid (AA) metabolism, and the folate pathway – the latter of which has a specific set of genes that regulate the cytoplasmic and mitochondrial arms (**Supplemental Figure 4A)**.

As above with eIF2α western blotting, we first chose to investigate the expression of these genes in the 30-week post-TAM hearts, as these animals had the most severe phenotype. These data demonstrate striking and robust differences in the expression of specific genes from these ISR regulated pathways, including *Atf4* and *Mthfd2* (**Supplemental Figures 4B-C)**, but also those related to AA synthesis including *Asns* (asparagine synthesis), *Psat* (serine synthesis) and *Pycr2* (proline synthesis) (**Figure 4E)**. Moreover, there was a clear difference in folate pathway genes, demonstrating a major upregulation of genes associated with the mitochondrial arm of the folate pathway (*Mthfd2*, *Shmt2*, *Aldh1l2*) (**Figure 4F)**, but an almost opposite effect on genes that regulate the cytoplasmic arm (*Mthfd1*, *Shmt1*, *Aldh1l1*) (**Figure 4G)**. In fact, several of the cytoplasmic folate pathway genes are significantly downregulated at this end-stage time point. Given these differences in the late-stage phenotype, we wanted to understand if these same differences in cytoplasmic versus mitochondrial folate gene activations were also observed in the early time points - prior to major functional defects in the heart. Indeed, similarly to the data from the 30-week mice, we show that 1C-met and AA synthesis genes are already significantly altered at the 20-week time point, and continue to be modulated as the phenotype progresses (**Figure 4H)**. It also appears that the mitochondrial arm of the folate pathway is specifically regulated in the mutant hearts at 20, 24 and 28 weeks post-TAM (**Figure 4I)**, with little effect on the cytoplasmic arm (**Figure 4J)**. These data strongly suggest that one of the early molecular events that perpetuates mitochondrial dysfunction and cardiomyopathy in these animals, is a specific defect in mitochondrial folate metabolism and accompanying activation of the ISR.

Other genes well known to be activated by the ISR pathway include the secreted peptide hormones FGF21 and GDF15. Several studies in mice and humans have shown that plasma abundance of these peptides is increased in the setting of ISR activation, and that they have chronic and robust impacts on peripheral metabolism and energy balance^23–26^. FGF21 imparts these effects through increasing energy expenditure and promoting adipocyte browning^27,28^, whilst GDF15 acts on satiety to reduce food intake^23,29^. Together they act in concert to potently regulate whole-body energy balance, and explains why in conditions where these factors are chronically elevated (in either mice or humans), significant weight loss is also observed. With this in mind, we next undertook experiments to test if FGF21/GDF15 were altered in PolG^Mut^ mice. Indeed, qPCR data demonstrates that in the LV of mutant mice, there were striking increases in the expression of both *Fgf21* and *Gdf15* at 20 weeks post TAM (**Figure 4K)**. Interestingly, this 20-week time point appears to be the most differentially regulated for *Fgf21* (∼100 fold), after which expression begins to wane (down to ∼40 fold at 28 weeks). On the contrary, whilst *Gdf15* is also substantially increased at the 20-week time point (∼16 fold) (**Figure 4L)**, its’ expression continues to escalate through the progressing time points (up to ∼40 fold at 28 weeks). With such striking differences in the expression of these two genes in the heart, we next assessed the abundance of these secreted factors in the plasma. These data demonstrate an increase in GDF15 at 20, 24 and 30 weeks post-TAM, but no major differences in FGF21 plasma levels (**Figures 4M-N**). The reason for this discrepancy in the amount of change of these circulating factors in the plasma, likely relates to the absolute basal abundance. FGF21 is already in high abundance in the plasma (∼3500pg/mL) from tissues such as the liver, and therefore the contribution coming from the mutant hearts has little impact on the total plasma pool. However, GDF15 has a comparatively much lower overall abundance (∼75pg/mL), and therefore the amount being secreted by the mutant hearts has a major impact on the total plasma abundance. The GDF15 data, which serves as a plasma-based proxy for ISR activation, provide further evidence that the ISR is an early and potentially precipitating event in the development of this phenotype. These data also suggest that it is highly likely that with such robust and chronic elevations in GDF15 (and potentially other factors secreted from the mutant heart), that these would partly explain the metabolic impacts we observed in the mutant animals (i.e. loss of fat mass), that may also drive changes in other peripheral tissues.

In summary of these data, we demonstrate that cardiomyocyte specific impairment of PolG activity drives increased damage to mtDNA, which activates the ISR in cardiomyocytes and subsequently promotes a robust and highly specific change in the folate pathway – particularly the mitochondrial arm of this metabolic pathway. Such findings suggest that chronic ISR activation constantly draws on metabolites from the mitochondrial folate pathway, which are required for processes such as mtDNA synthesis, mitochondrial antioxidant defence, and mitochondrial protein translation. This constant need for metabolites from this pathway leads to a transcriptional feedback aimed at increasing the activity of the mitochondrial folate pathway, however the ultimate failure of this feedback presumably leads to chronic depletion of folate metabolites and contributes to mitochondrial dysfunction and cardiomyopathy.

### Peripheral Impacts of PolG induced Cardiomyopathy

As a final investigation in these mutant animals, we were interested in defining the changes that were occurring in peripheral tissues in response to cardiomyopathy induced by chronic ISR and mitochondrial dysfunction. As touched on above, this particular phenotype develops rapidly and severely, and does so with the heart releasing certain known (such as FGF21 and GDF15), and presumably unknown factors into the circulation. Whilst we suspect GDF15 has major impacts on energy metabolism in these animals, we wanted to demonstrate if direct changes were also occurring in other tissues including adipose tissue, skeletal muscle and liver. Additionally, it was also important to demonstrate that the ISR was not activated in these tissues, to further validate the “cardiac” specific nature of our model.

There was substantial fat mass loss in mutant animals in the last 4-6 weeks of the phenotype, with animals losing up to 60-70% of their starting white adipose tissue (WAT) weight (epididymal and subcutaneous) at the most end-stage timepoint (Supplemental Figure 1F). Therefore, we first focused on molecular pathways being impacted in WAT. This was performed using qPCR analysis for key transcriptional and metabolic genes, which demonstrates rewiring of a number of lipid metabolism and adipogenic developmental gene programs (consistent with adipose tissue remodelling) (**Figure 5A**). Western blotting for key readouts of the ISR in adipose tissue demonstrated no change in phosphorylation (Ser51) of eIF2α (**Figure 5B** and **Supplemental Figure 5D**), nor was there any change in expression of ISR related genes by qPCR (**Figure 5C**), indicating that adipose had no signs of activation of this pathway.

**Figure 5:**
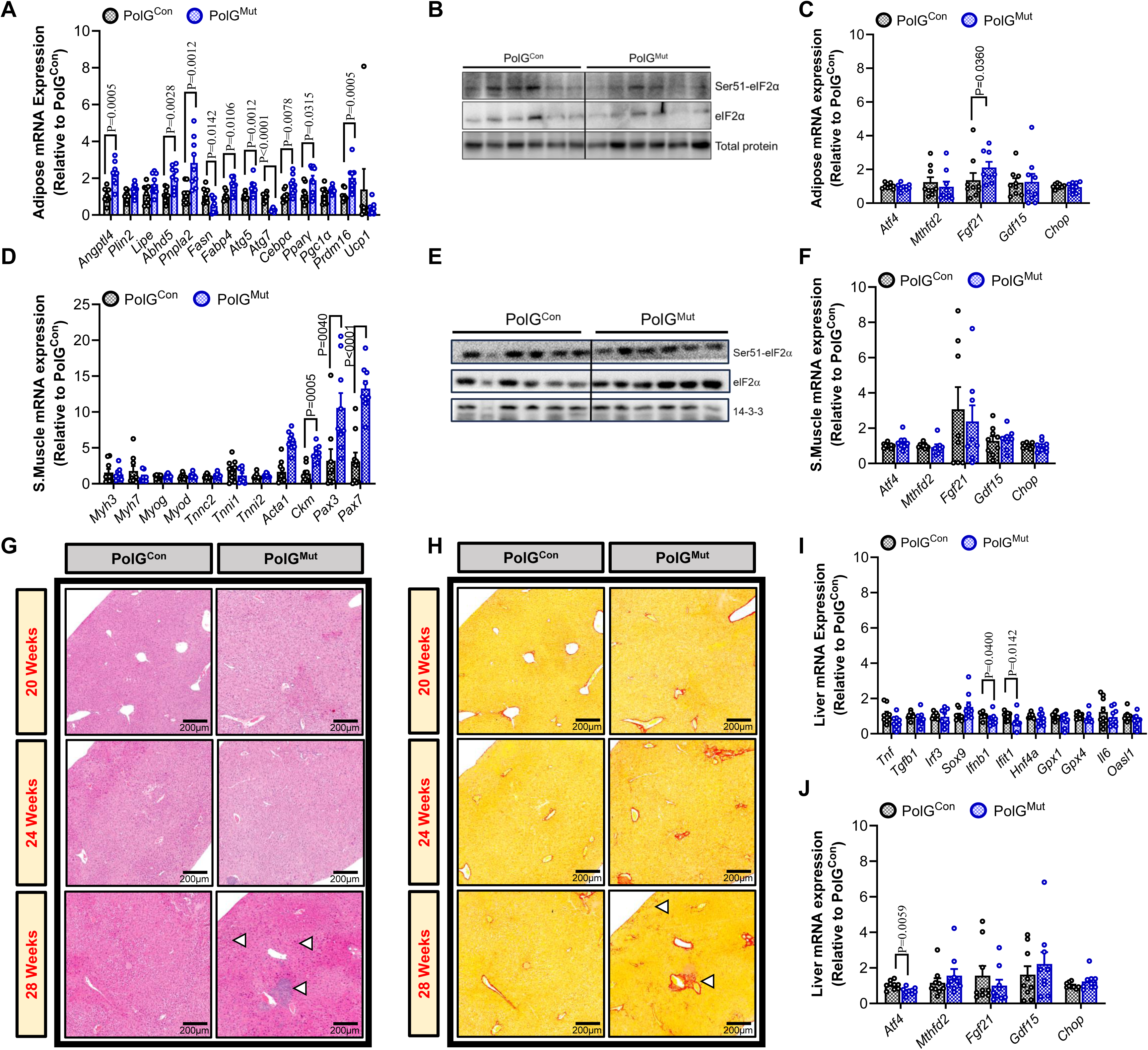
Activation of the integrated stress response specifically in cardiomyocytes contributes to peripheral liver damage. **(A)** Gene expression (qPCR) in subcutaneous white adipose tissue (inguinal) from PolG^Mut^ and PolG^Con^ mice at 28 weeks post-Tam (n=9/group). **(B)** Immunoblot of phosphorylated and total eIF2α (n=6/group) and **(C)** expression of ISR genes (n=9/group) in inguinal adipose from PolG^Mut^ and PolG^Con^ mice at 28 weeks post-Tam. **(D)** Gene expression (qPCR) in skeletal muscle (tibialis anterior) from PolG^Mut^ and PolG^Con^ mice at 28 weeks post-Tam (n=9/group). **(E)** Immunoblot of phosphorylated and total eIF2α (n=6/group) and **(F)** expression of ISR genes (n=9/group) in tibialis anterior muscle from PolG^Mut^ and PolG^Con^ mice at 28 weeks post-Tam. Representative images of PolG^Mut^ and PolG^Con^ mice liver sections stained with **(G)** haematoxylin and eosin stain (white arrows show regions of inflammation/necrosis) and **(H)** picrosirius red at 20, 24 and 28 weeks post-Tam (white arrows show strong areas of fibrosis). Expression (qPCR) of **(I)** immune genes and **(J)** ISR genes in liver from PolG^Mut^ and PolG^Con^ mice at 28 weeks post-Tam (n=9/group). Data are presented as mean ± SEM, with P-value between control and mutant biological replicates determined by unpaired welch t-test **(J)** or Mann-Whitney test **(A, C, D, F, I)**. Source data for these figures are provided in the Source Data file.

Next, we sought to investigate the impacts of the phenotype on skeletal muscle function. The reasons for this were two-fold; firstly, end stage cardiomyopathy is often accompanied by muscle wasting, and secondly, our previous skeletal muscle specific mutant model demonstrated a strong signal for muscle degeneration and regeneration occurring^11^. Therefore, in quadricep muscle of mice from the end-stage time point in the current study (30 weeks post-TAM), we investigated pathways relevant to ISR and to muscle remodelling. Using qPCR, we demonstrate that there was some minor change in a handful of genes relating to muscle remodelling of skeletal muscle, indicating that there was perhaps a modest impact of cardiomyopathy on muscle integrity and mobilisation of satellite cells in muscle tissue, as evidenced by increased expression of intermediate structural proteins (*Acta1* and *Ckm*), and Pax transcription factors (*Pax3* and *Pax7*), respectively (**Figure 5D**). However, there was no change in the phosphorylation (Ser51) of eIF2α in quad muscle (**Figure 5E** and **Supplemental Figure 5E**), nor any change in genes associated with ISR activation (**Figure 5F**), confirming that there was no evidence for activation of the ISR in skeletal muscle, and that these muscle remodelling effects were likely secondary complications associated with cardiomyopathy.

Lastly, we investigated the impacts of the PolG cardiomyopathy model on liver health. Here, there was a striking phenotype that was both unexpected, yet highly relevant to the clinical setting. In the 30-weeks post-TAM time point, we noted the development of a nutmeg-like appearance in the livers of the majority of mutant mice, which was accompanied with what appeared to be localised necrosis and cirrhosis (**Supplemental Figure 5A**). Mild evidence of this phenotype was also observed in the 28-week livers (**Supplemental Figure 5B**). This phenotype is known as cardiac hepatopathy (CH), and has been identified in up to 80% of human patients with HF^30–32^. Histological examination of these livers also showed characteristic features of CH, including localised regions of necrosis and inflammatory infiltration (**Figure 5G**, white arrows), along with an increase in fibrosis (**Figure 5H**, white arrows). Using qPCR, we demonstrate that there is only a minor change in specific gene sets associated with metabolism and remodelling pathways, with the most significant changes observed in lipid metabolism genes (**Figure 5K** and **Supplemental Figure 5C**).

In humans, CH is thought to be driven in part by reductions in blood flow, but also due to local and extrinsic factors released into the circulation that drive subsequent phenotypes. This includes FGF21 and GDF15, both of which are being modulated in the hearts of our mutant mice. CH appears to be a rapidly progressing pathology in our model, as we observe only minor features of the condition at the 28 week time point, with negligible signs at 24 weeks (**Supplemental Figure 5B**). To our knowledge this is the first description of a mouse model with cardiomyopathy that presents with this phenotype, and therefore represents a model of significant relevance to studying hepatic complications associated with cardiomyopathy and HF. We believe that CH in this model is specific to cardiomyopathy induced by mitochondrial dysfunction and activation of the ISR, and not to generalised impacts on blood flow, as we have not observed striking liver phenotypes previously in other preclinical models of HF across our broader group including ischemia reperfusion, permanent coronary ligation, transverse aortic constriction or genetic models (PI3K-Tg, Mst1-Tg). Thus, we propose that this model will be an important tool moving forward to aid in discovery of pathways that both promote and progress the development of CH, with opportunities to identify new diagnostics and therapeutics to manage this escalating clinical problem.

In summary, we present a novel mouse model of cardiomyopathy that is induced by impairing the activity of the mtDNA modulating enzyme Polymerase gamma (PolG), specifically in cardiomyocytes. This PolG impairment drives increases in mtDNA damage and chronically activates the ISR. Our data suggests that this chronic activation of the ISR likely depletes cardiomyocytes of critical intermediates in the mitochondrial folate pathway, culminating in mitochondrial dysfunction and ultimately cardiomyopathy. As part of the chronic activation of the ISR, factors are secreted from the injured heart that pass into circulation and have major impacts on peripheral tissues including the liver, that likely drive conditions such as cardiac hepatopathy. This model thus represents a major advance in our understanding of the mechanisms that drive cardiomyopathy and its complications, highlighting potential new avenues for therapy including modalities that replenish levels of mitochondrial folates, and those that dampen chronic activation of the ISR.

## DISCUSSION

The cardiomyocyte specific PolG mutant model presented in this study, provides major insights into new and emerging mitochondrial stress mechanisms that drive degenerative disease. The model circumvents systemic confounders that exist in other mouse models of PolG related disease, and allows for temporal and tissue specific investigation into the signalling cascades activated by mtDNA damage in cardiomyocytes. This includes clear evidence for early and sustained activation of the integrated stress response (ISR), through to severe structural and functional cardiac defects. A highly relevant pathway of interest that has emerged from this work, is the robust and specific impacts on mitochondrial folate metabolism. This is consistent with our previous work in skeletal muscle, and thus we propose that this is an early and sustained defect that perpetuates and exacerbates degenerative disease – perhaps exclusively in post-mitotic tissues.

Whilst this model provides what might be perceived as a small advance in the toolbox of available models to study PolG specific biology, the inducible and temporal cardiac specific studies that are possible using this model, have allowed us unprecedented capacity to tease out the intricate timing of the various pathological insults that ultimately drive cardiac failure in the setting of mtDNA damage. A major contributor to this pathology is the chronic activation of the ISR and the development of cardiac hepatopathy, the latter of which has never before been reported in PolG related models. We show that the ISR is likely the earliest detectable pathway that is activated in response to an increasing burden of mtDNA damage in cardiomyocytes, and the pathway remains chronically activated until the heart ultimately fails, leading to death. These molecular events occur before overt reductions in cardiac function, demonstrating that ISR activation is not merely a secondary response to heart failure, but likely plays a causal role in disease progression. This evidence supports the notion that mitochondrial triggered ISR initiates maladaptive cellular processes, independent to or preceding ATP depletion, that ultimately promote the development of HF.

Whilst the ISR is well described in its important roles in resolving acute cellular insults, and indeed is necessary for cell survival in some circumstances, it is only more recently that chronic and sustained ISR activation has been studied in its role as a driver of pathology. One such study published recently by Takaoka and colleagues^33^, engaged a novel genetic mouse model that specifically led to chronic activation of the ISR in the heart, independent to any effects relating to mitochondrial function. This was achieved by deleting GADD34 (*Ppp1r15a*), a gene that encodes a phosphatase that deactivates eIF2α. Thus, by deleting GADD34 in the heart, when they exposed KO animals to irradiation stress, the ISR was chronically activated in the heart and KO animals developed severe HF. Importantly, control animals that were replete of GADD34 did not develop HF following irradiation exposure, nor did KO animals develop heart failure in the absence of irradiation therapy. This paper therefore supports the concept presented in our current study, that chronic activation of the ISR in the heart is sufficient to drive cardiomyopathy.

Whilst our data provide important fundamental insights, there are still questions that remain unresolved with regards to the underlying molecular drivers. For example, in cells that are accumulating mtDNA damage, what is the “switch” that ultimately leads to activation of eIF2α? Moreover, how does chronic activation of the ISR ultimately drive pathology? We provide insight into both of these questions in our current and prior work. Firstly, we have previously shown using the same PolG floxed model, that mtDNA damage specifically in skeletal muscle (using an inducible ACTA1-cre model), activates the ISR and leads to maladaptive responses in skeletal muscle that drive muscle degeneration and wasting^11^. With regards to the likely upstream kinase to eIF2α, there are currently four known (PKR, PERK, GCN2 and HRI)^34^. Previous studies have investigated the potential upstream kinase in cell models such as cultured skeletal myotubes. These studies identified GCN2 as a kinase that was activated in response to Complex I inhibition, increased ROS production and thus depletion of asparagine^35^. In our skeletal muscle mouse model, we provided evidence that HRI was the likely upstream kinase. HRI resides in the cytosol, but has been shown to be activated by a cleaved variant of the protein DELE1 (sDELE1), which is converted in the mitochondria by the protease OMA1 and exported into the cytosol where it binds and activates HRI^9,36^. We thus predict that HRI is the likely pathway that would be active here in our cardiomyocyte specific model.

With regards to how chronic ISR drives degenerative pathology, we suspect that a sustained and ultimately untenable strain on metabolite flux through the mitochondrial arm of the folate cycle, is a strong contributing factor. This is evidenced by the clear and robust effort in mutant cells to increase expression of enzymatic machinery that would generate more substrates via the mitochondrial folate arm of the pathway (i.e. *Mthfd2*, *Shmt2*, *Aldh1l2*). However, this appears to be inadequate to sustain supply sufficiently and thus the 1C-metabolism network fails, leading to maladaptive pathologies. It is unclear why the increased expression of these enzymes is insufficient to keep up with demand, and it is possible that there are other potentially unknown post-translational blocks in one or more of these biochemical pathways that are responsible for this defect. This is even more intriguing when considering that the expression of genes from the cytoplasmic arm of the folate pathway (i.e. *Mthfd1*, *Shmt1*, *Aldh1l1*), appear to decrease in the mutant cells. This is either a further effort to shunt more folates through the mitochondria, or perhaps in response to an accumulation of a cytoplasmic folate derivative that arises due to failure of the mitochondrial arm. In either case, these data provide strong support for further work in this area to identify these specific defects, and to also provide opportunities to remedy these defects through either genetic or pharmacological means. Such interventions may reduce activation of the ISR and subsequent degenerative pathologies. In the context of the current study, these results highlight the ISR as a potential therapeutic target for mitochondrial cardiomyopathies or other diseases with underlying chronic ISR activation, where pharmacologic modulation of the ISR or mitochondrial folate pathways could offer new approaches to mitigate cardiac remodelling and heart failure.

Another intriguing finding from our study was the observation of cardiac hepatopathy in the cardiomyocyte specific mutants. This phenotype developed in the later stage animals (from 28 weeks onwards) and was overt in at least 50% of end stage animals (30 weeks post-TAM). Cardiac hepatopathy, or “nutmeg” liver^37^, has been loosely categorised into two main forms; congestive hepatopathy and Acute Cardiogenic Liver Injury (ACLI). ACLIs, such as ischemic hepatitis, are caused by hypoperfusion-induced hypoxia and congestion in the liver, whereas congestive hepatopathy is linked to elevated right sided and central venous pressures resulting in passive congestion within the liver^38^. However, clinical and indeed preclinical presentation of congestive hepatopathy is highly variable, yet reduced blood flow and congestion occurs in a high proportion of individuals with HF. This strongly suggests that additional contributing factors beyond just insufficient blood flow, congestion and hypoxia, play a role in the development of cardiac hepatopathy (94). This may be due, in part, to the release of circulating factors from the failing heart, such as FGF21, GDF15, tumour necrosis factor alpha (TNF-α) and myosin heavy chain 6 (Myh6), which have previously been implicated in liver dysfunction (94, 95). Indeed, GDF15 is currently being explored as a biomarker of HF, as it is secreted by the heart during the cardiac remodelling process (96, 97). Interestingly, clearance of GDF15 from the circulation using antibodies, was sufficient to improve HF in a mouse model of ISR driven HF^33^, and thus cardiac released factors are of high relevance to both the progression of HF itself, and its complications.

With this in mind, we propose that the CH observed in the current model is specific to HF associated with ISR, and not just to generalised reductions in blood flow that are associated with the vast majority of HF models. This is supported by the fact that we, and presumably others (due to a lack of reports on CH), have not observed overt CH previously in other models of HF across our broader group including coronary ligation induced ischemia^39^, transverse aortic constriction^40^ and genetic models^41,42^. Thus, we propose that this model will be an important tool moving forward to aid in discovery of pathways that both promote and progress the development of CH, with opportunities to identify new diagnostics and therapeutics to aid in managing this escalating clinical problem.

Some limitations exist in the current work further to the questions already raised above, including that this work was only performed in male mice, so it is unknown whether the phenotype progresses similarly in females. Moreover, we have only performed the deletion in an inducible setting (starting from 8 weeks of age), which although is highly desirable in terms of eliminating potential developmental and compensatory mechanisms, cannot provide insights into how the truncation impacts heart development from conception. Finally, whilst our data strongly implicates both the ISR and mitochondrial folate pathways in exacerbating the phenotype in these mutants, we have not shown any data on the impact of manipulating these pathways on disease outcomes. Whilst this is obviously outside the scope of the current work, it is nonetheless a critical set of experiments to perform in order to convincingly demonstrate the therapeutic relevance of these pathways in the development of HF.

In summary, our cardiomyocyte-specific PolG^Mut^ model establishes a direct mechanistic link between mtDNA damage, ISR activation, altered folate metabolism and the development of severe cardiomyopathy and CH. This model provides a unique opportunity to further dissect ISR signalling in the heart and to identify therapeutic strategies aimed at halting or reversing ISR-driven cardiac pathology, as well being a new and unique model for the study of cardiac hepatopathy.

## METHODS

### Animals

All animal experiments were approved by the Alfred Research Alliance (ARA) Animal Ethics committee (E/8347/2024/B) and performed in accordance with the ethical guidelines set out by the National Health and Medical Research Council (NHMRC) of Australia. Floxed PolG mutator mice (Polg^fl/fl^) were generated in collaboration with the Monash Genome Modification Platform as previously described^11^. Recombination was achieved using the Cre-Lox system by crossing the Polg^fl/fl^ mice with MHCα-CreERT2 mice (C57BL/6J background, Jackson Laboratories, #005657) to generate Polg^fl/fl^-MHCα-CreERT2^−/−^ (Polg^fl/fl^) and Polg^fl/fl^ MHCα-CreERT2^+/−^ (Polg^fl/fl-Cre^) animals. For the first cohort, male mice were bred and genotyped, before being randomly allocated into two separate groups each, totalling four groups altogether. Mice were aged to 6–8 weeks before one group of each genotype received a single intraperitoneal injection of Tamoxifen (40mg/kg) in sunflower oil, whilst the other received sunflower oil alone. This resulted in four different groups: Polg^fl/fl+OIL^, Polg^fl/fl-Cre+OIL^, Polg^fl/fl+TAM^ (PolG^cont^), Polg^fl/fl-Cre+TAM^ (PolG^mut^). Mice were housed at 22°C on a 12 h light/dark cycle with access to food (chow: Specialty feeds, Australia) and water *ad libitum* with cages changed weekly. These cohorts were aged up to 30-weeks post-tamoxifen. For subsequent cohorts, only PolG^cont^ and PolG^mut^ mice were generated as above and studied. Three different cohorts were generated and followed until end point, which was 20, 24 and 28 weeks post-tamoxifen. At the end of the studies, mice were fasted for 4–6 hrs and then anesthetized with a lethal dose of ketamine/xylazine before blood and tissues were collected, weighed and frozen for subsequent analysis.

### EchoMRI

Body composition analysis, including lean mass (LM) and fat mass (FM), was analysed in living mice using the 4-in-1 NMR Body Composition Analyzer for Live Small Animals (EchoMRI LLC, Houston, TX, USA). Analysis was performed at designated periods throughout the study as previously described^43–45^.

### Mitochondrial (mt)DNA to nuclear (n)DNA ratio

Mitochondrial content was determined by qPCR using a ratio of mtDNA to nDNA, as previously described ^46^. Briefly, a ∼50mg piece of the left ventricle was homogenised in digestion buffer (100 mM NaCl, 10 mM Tris-HCl, 25 mM EDTA, 0.5% SDS, pH 8.0) and then incubated in Proteinase K (250U/mL) for 1 h at 55°C. Following this, total DNA was isolated using the phenol-chloroform extraction method. A qPCR reaction was then performed on 2ng of total DNA using a primer set that amplifies the mitochondrial gene *Mtco3*, and the nuclear gene *Sdha* (see **Table 1** for primer sequences). Estimated abundance of each gene was used to generate a ratio of mitochondrial to nuclear DNA (mtDNA/nDNA), and this ratio was compared between genotypes.

### Blood Pressure measurement in conscious mice (tail cuff)

Blood pressure was measured in conscious mice using tail cuffs with a Kent Scientific CODA non-invasive blood pressure system. Measurements were performed in PolG^Con^ and PolG^Mut^ mice at 20, 24, and 28 weeks post-Tam. Prior to recordings, mice were acclimated to the procedure over two consecutive days by placing each animal in the restraint holder on the heated platform for 5 minutes per day. On the third day, blood pressure data was collected. Blood pressure was measured by restraining mice in the clear restraint tube placed on the heated platform with the sensor cuffs placed over the tail. Ten consecutive blood pressure measurements were recorded per mouse. The two highest and two lowest values were excluded, and the median of the remaining six measurements was used for analysis. Both systolic and diastolic pressures were recorded.

### Quantitative PCR (qPCR)

RNA was isolated from tissues using RNAzol reagent and isopropanol precipitation. cDNA was generated from RNA using MMLV reverse transcriptase (Invitrogen) according to the manufacturer’s instructions. qPCR was performed on 10ng of cDNA using the SYBR-green method on a QuantStudio 7 Flex Real-Time PCR System, using primer sets outlined in **Table 1**. Primers were designed to span exon-exon junctions where possible, and were tested for specificity using BLAST (Basic Local Alignment Search Tool; National Centre for Biotechnology Information). Amplification of a single amplicon was estimated from melt curve analysis, ensuring an expected temperature dissociation profile was observed. Quantification of a given gene was determined by the relative mRNA level compared with control using the delta-CT method, which was calculated after normalisation to one of two housekeeping genes; *Ppia* or *Rplp0*.

### Mitochondrial Isolation

For isolation of mitochondria, a piece of left ventricle was incubated and minced in ice-cold isolation buffer A (220mM Mannitol, 70mM Sucrose, 20mM HEPES, 2mM Tris-HCl, 1mM EDTA/EGTA, pH 7.2) + 0.4% (w/v) fatty acid free BSA and then homogenized with a glass dounce homogenizer. The homogenate was centrifuged at 650 g for 5min at 4°C and the pellet was discarded, and the supernatant transferred to a fresh tube. The supernatant was repeatedly centrifuged at 650 g for 5 minutes until very little material was pelleted. The supernatant was then transferred to a high-speed centrifuge tube and centrifuged at 10,000 g for 5 minutes and the supernatant was discarded, and the crude mitochondrial pellet resuspended in isolation buffer A. Mitochondria were re-pelleted by centrifuging at 10,000 g for 5 minutes, and the supernatant was discarded, and the pellet resuspended in 1ml of isolation buffer B (220mM Mannitol, 70mM Sucrose, 10mM Tris-HCl, 1mM EDTA, pH 7.2). The final pellet was collected by centrifuging at 10,000 g for 5 minutes, discarding the supernatant and resuspending the pellet in either 1 x PBS for proteomic analysis or isolation buffer B.

### Mitochondrial Sequencing

1μg of genomic DNA was precipitated from 20-week and 28-week post-tamoxifen PolG mutant and control hearts. Mitochondrial DNA was sequenced by CD Genomics (New York, USA).

### Droplet Digital PCR (ddPCR)

The RMCA was performed similarly to that previously described^47^. Briefly, mitochondria were isolated from skeletal muscle as described above. Total DNA was then extracted using the phenol chloroform precipitation method. 50ng of mtDNA was digested for 6 hours with *TaqI* enzyme, with fresh enzyme added every hour before a final overnight incubation followed by inactivation at 80°C. 5ng of digested mtDNA was directly analysed on a BioRad QX200 Droplet Digital PCR System using primers that span the *TaqI* sensitive 634 site, where as 0.05ng of digested mtDNA was used for detection of the control, *TaqI* insensitive site (mtCont) (see **Table 1** for primer sequences). Assays were performed using EvaGreen reaction mix according to the manufacturer’s instructions using the following conditions: dissociation (95°C for 30s), annealing (61.8°C for 1min) and extension (72°C for 1.5min) for 50 cycles. Only samples where >10,000 droplets were quantified and included in subsequent analysis.

### Conventional PCR for mtDNA Deletions

To visualise mtDNA deletions we performed conventional PCR on isolated mtDNA from left ventricle, as previously described^11^. Briefly, 10ng of mtDNA precipitated from left ventricle mitochondria (previously treated with DNAse I), was amplified using Phusion Polymerase according to the manufacturer’s instructions, using the “mtDNA deletion” primers outlined in **Table 1** (which flank the major arc of the mtDNA molecule; ∼10kb). The following conditions were used in the amplification reaction: dissociation (95°C for 30s), annealing (60°C for 30sec) and extension (72°C for 5min) for 40 cycles.

### SDS-PAGE and immunoblot

Tissues were lysed in radio-immunoprecipitation assay (RIPA) buffer supplemented with protease and phosphatase inhibitors. Matched protein quantities were separated by SDS-PAGE and transferred to PVDF membranes. Membranes were blocked in 3% skim milk for 2 h and then incubated with primary antibody (see **Table 2** for details) overnight at 4°C. After incubation with primary antibodies, membranes were washed and probed with their respective HRP-conjugate secondary anti-mouse or anti-rabbit (BioRad) antibodies in 3% skim milk for 2h at room temperature, then visualised with enhanced chemiluminescent substrate (Pierce). Approximated molecular weights of proteins were determined from a co-resolved molecular weight standard (BioRad, #1610374). Image Lab Software (BioRad) was used to perform densitometry analyses, and all quantification results were normalized to their respective loading control or total protein.

### Plasma FGF21 & GDF15

Commercial ELISA kits for FGF21 (R&D Systems, #MF2100) and GDF15 (R&D Systems, #MGD150) were used to measure respective plasma concentration as per manufacturer’s instructions.

### Proteomics

#### Proteomic sample preparation

Mouse heart tissue (20-, 24- & 28-week PolG^Con^/PolG^Mut^, n=5/group/time-point) were lysed on ice (8M urea in 50 mM HEPES pH 8.0 with Halt™ Protease/Phosphatase Inhibitor Cocktail, #78442, Thermo Fisher Scientific) and extracted by rapid pulse tip-probe sonication on ice. Heart tissue lysates were quantified (microBCA, #23235, Thermo Fisher Scientific), normalised, and a modified sera□bead workflow employed as described^48,49^. Briefly, samples were solubilised in 2% v/v sodium dodecyl sulphate (SDS) incubated for 5 min at 95°C, reduced (10mM dithiothreitol (DTT) for 1 hr at 25°C) and alkylated (20mM iodoacetamide (IAA) for 30 min in dark at 25 °C). Samples were quenched with 10 mM DTT and combined with Sera-Mag Speed Beads mixtures (#45152105050250 and #65152105050250, GE LifeScience) at a 10:1 beads-to-protein ratio as described^49^. Lysates were reconstituted in 50mM TEAB pH 8 and enzymatically digested with trypsin (V5113, Promega, 1:50) and Lys-C (121-05063, FUJIFILM Wako Pure Chemical Corporation, 1:100 enzyme-to-substrate ratio) overnight at 37°C with shaking. Peptides were acidified, lyophilised and quantified as described^49^ before analyses.

#### NanoLC and Mass Spectrometry

Peptides were analysed on a Dionex UltiMate NCS□3500RS nanoUHPLC coupled to a Q□Exactive HF□X hybrid quadrupole□Orbitrap mass spectrometer equipped with nanospray ion source in positive, data-independent acquisition mode as described^48,49^. Peptides were loaded (Acclaim PepMap100 C18 5 µm beads with 100 Å pore□size, Thermo Fisher Scientific) and separated (1.9 µm particle size C18, 0.075 × 250 mm, Nikkyo Technos Co. Ltd) with a gradient of 2–28% acetonitrile containing 0.1% formic acid over 45 mins followed by 28-80% from 45-47 min for total runtime of 56 min at 300 nl min-1 at 55 °C (butterfly portfolio heater, Phoenix S&T). Full scan MS were performed in the m/z range of 350 to 1100 m/z with a 60,000 resolution, using an automatic gain control (AGC) of 3x10^6, maximum injection time of 50 ms and 1 microscan. MS2 was set to 15,000 resolution, 1e6 AGC target and the first fixed mass set to 120 m/z. Default charge state set to 2 and recorded in centroid mode^49^. Data were acquired using Xcalibur software v4.5 (Thermo Fisher Scientific).

MS-based proteomics data is deposited to the ProteomeXchange Consortium via the MassIVE partner repository and available via MassIVE with identifier MSV000100303.

#### MS data processing and analysis

For global proteome analysis, identification and quantification of peptides and proteins was performed using DIA-NN neural network and interference correction (v1.9) in library-free mode searched against *Mus musculus* (mouse) reference proteome (55,398)^50^. Spectral libraries were predicted using the deep learning algorithm employed in DIA-NN with Trypsin/P, allowing up to 1 missed cleavage. The precursor charge range was set to 1-4, and the m/z precursor range was set to 300-1800 for peptides consisting of 7-30 amino acids with N-term methionine excision and cysteine carbamidomethylation enabled as a fixed modification with 0 maximum number of variable modifications. The mass spectra were analysed using default settings with a false discovery rate (FDR) of 1% for precursor identifications and match between runs (MBR) enabled.

#### Proteome data analysis and informatics

Data analysis and summary values were performed using Perseus (v2.0.11.0) of the MaxQuant software package. Data quality filters were applied to global datasets with a minimum of 60% protein identification rate in at least one sample group before proceeding to downstream statistical analysis. Protein intensities were log2 transformed and normalized using quantile normalization. Principal component analysis (PCA) was performed with missing values imputed from normal distribution (width 0.3, downshift 1.8); however, imputed values were excluded from subsequent statistical analyses. For differential analysis, one-way ANOVA and Student’s t-tests were applied with permutation-based false discovery rate (FDR) correction, and hierarchical clustering was performed using Euclidean distance and average linkage clustering. Data visualization was performed using Perseus, R (ggplot2) package, Microsoft Excel and GraphPad Prism (v10) to generate boxplots, volcano plots, heatmaps, and bar charts. Pathway analysis was performed in ShinyGO 0.85.1 (South Dakota State University, USA)^51^ analysing the top 15 up and downregulated KEGG^52^ pathways (minimum pathway size of 2 with FDR cutoff at 0.15). Significant pathways were sorted by pathway enrichment.

### Histology

Hearts were fixed in formalin and mounted in paraffin before 4µm sections were cut on a Leica microtome. All sections were stained with Pico Sirius Red (PSR) or haematoxylin and eosin stain (H&E), and slide images were captured using an Zeiss Axioscan 7(Zeiss, Germany) and viewed in the accompanying software ZEN (Zeiss, Germany) & QuPath (United Kingdom).

### Cardiac Echocardiography

Transthoracic echocardiography was performed using a Vevo 2100 Ultrasound Machine (VisualSonics) with a 40MHz transducer in anaesthetized mice (1.5-2% isoflurane, inhalation) to assess cardiac structure and function. All data were collected blinded and validated by two independent researchers. Measures included: heart rate, end systolic volume, ejection fraction, cardiac output, stroke volume, global longitudinal strain and fractional shortening.

### Statistical analyses

All data were expressed as mean ± standard error of the mean (SEM) unless otherwise stated. Normality was checked using Shapiro-Wilk normality tests. All statistical analyses in animal studies were analysed by repeated measures two-way ANOVA (for normally distributed data), other than where explicitly detailed. Tissue analysis experiments were analysed by ANOVA with post hoc testing (Fishers LSD) where appropriate, or unpaired welch t test or Mann-Whitney test unless otherwise stated. Analyses were performed using PRISM9 software and a *p* value of *p* < 0.05 was considered statistically significant.

### Data inclusion and exclusion criteria

For animal experiments, phenotyping data points were excluded using the following pre-determined criteria: if the animal was unwell at the time of analysis, there were technical issues identified, or data points were identified as outliers using Tukey’s Outlier Detection Method (1.5 IQR < Q1 or 1.5 IQR > Q3). If repeated data points from the same mouse/sample failed QC based on predetermined criteria or several data points were outliers as per Tukey’s rule, the entire animal was excluded from that given analysis. For *in vivo* and *in vitro* tissue and molecular analyses, data points were only excluded if there was a technical failure (i.e. failed amplification in qPCR, failed injection in mass spectrometer) or the value was biologically improbable. This was performed in a blinded fashion (i.e., on grouped datasets before genotypes were known).

## SUPPLEMENTAL FIGURE LEGENDS

**Supplementary Figure 1:**
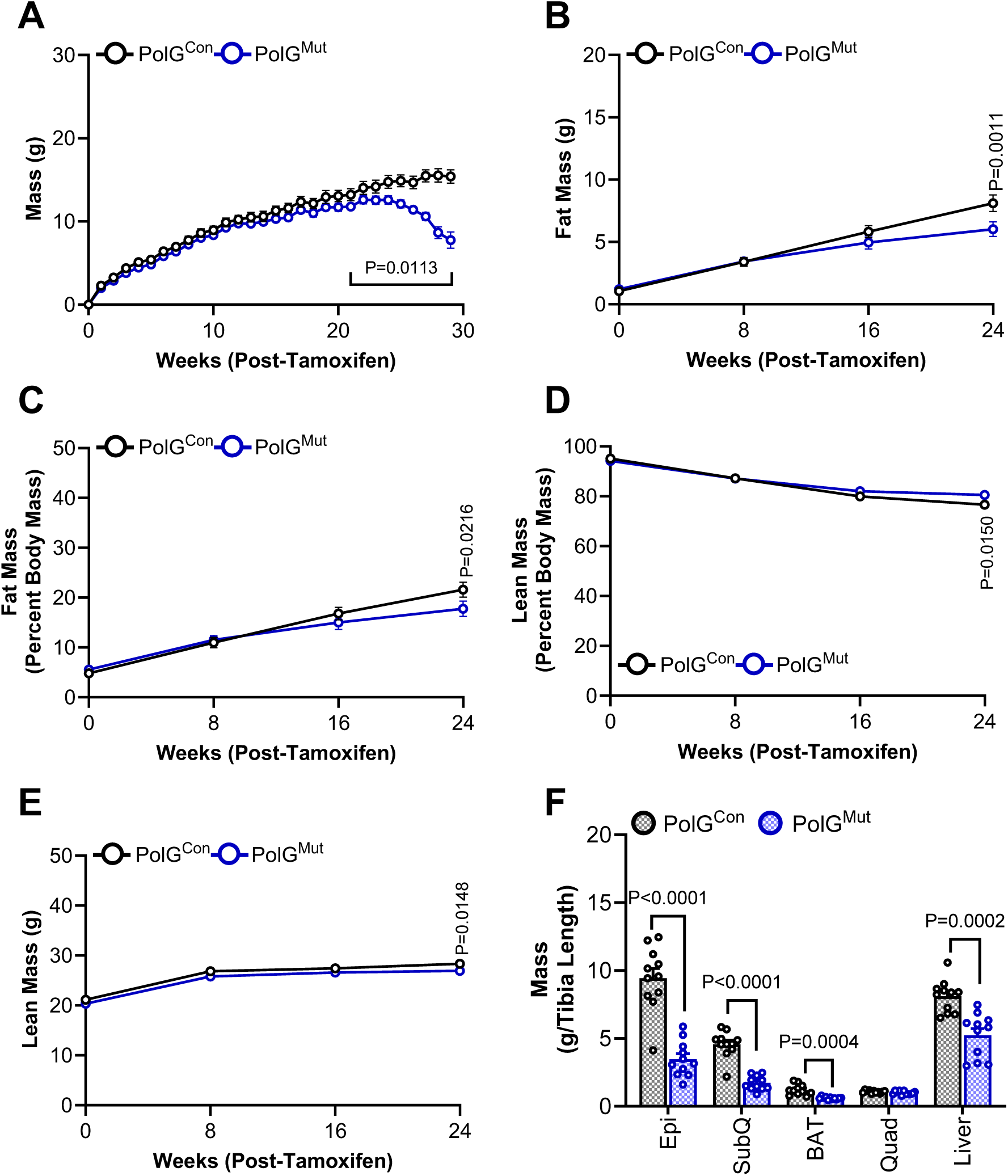
Mutation to PolG gene impacts body composition. **(A)** Weight gain from 0-30 weeks post-Tam in PolG^Mut^ (n=15) and PolG^Con^ (n=11) mice. Body composition of PolG^Mut^ (n=15) and PolG^Con^ (n=11) mice including **(B)** fat mass, **(C)** fat mass as a percentage of total body weight, **(D)** lean mass as a percentage of total body weight lean mass, and **(E)** lean mass in grams (g). **(F)** wet tissue weights of epididymal adipose(epi), inguinal adipose (SubQ), brown adipose (BAT), quadricep (Quad) and liver excised from PolG^Mut^ and PolG^Con^ mice (n=11/group). Data are presented as mean ± SEM, with P-value between control and mutant biological replicates determined by two-way ANOVA (mixed model) with correction for multiple comparisons **(A-E)**, or unpaired welch t-test **(F)**. Source data for these figures are provided in the Source Data file.

**Supplementary Figure 2:**
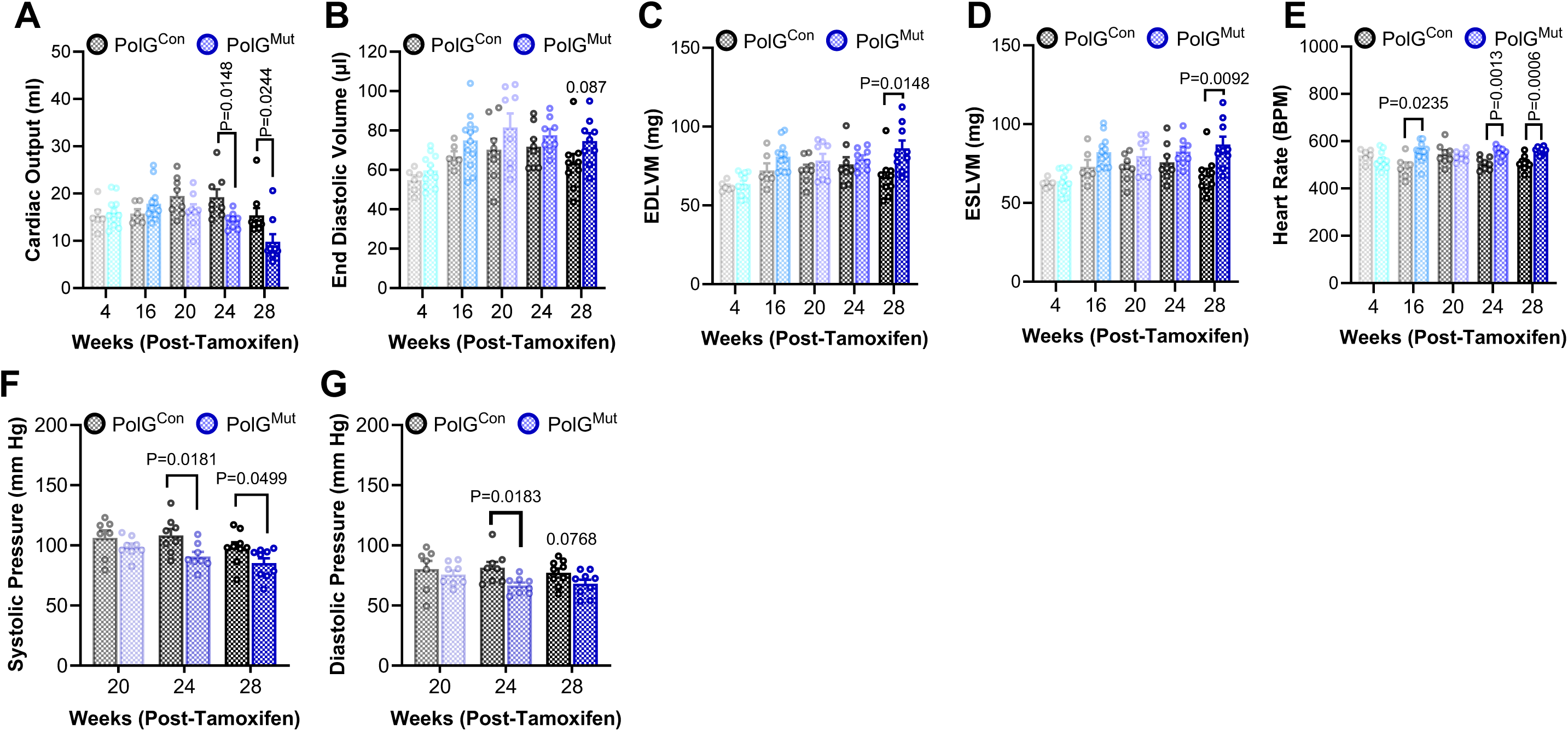
Progressive accumulation of cardiomyocyte mtDNA damage and cardiac dysfunction in PolG^Mut^ mice. Cardiac function, including **(A)** Cardiac output **(B)** end diastolic volume, **(C)** end diastolic left ventricular mass (EDLVM), **(D)** end systolic left ventricular mass (ESLVM), and **(E)** heart rate was assessed by echocardiography at 4 (PolG^Mut^ n=11 and PolG^Con^ n=6), 16 (PolG^Mut^ n=12 and PolG^Con^ n=6), 20 (n=8/group), 24 (n=8/group) and 28 (n=9/group) weeks post-Tam in PolG^Mut^ and PolG^Con^ mice. **(F)** Systolic and **(G)** Diastolic blood pressure measured by tail cuff in PolG^Mut^ and PolG^Con^ mice at 20 (PolG^Mut^ n=8, PolG^Con^ n=7), 24 (n=8/group) and 28 (n=9/group) weeks post-Tam. Data are presented as mean ± SEM, with P-value between control and mutant biological replicates determined by unpaired welch t-test **(A)** or Mann-Whitney test **(B-G)**. Source data for these figures are provided in the Source Data file.

**Supplementary Figure 3:**
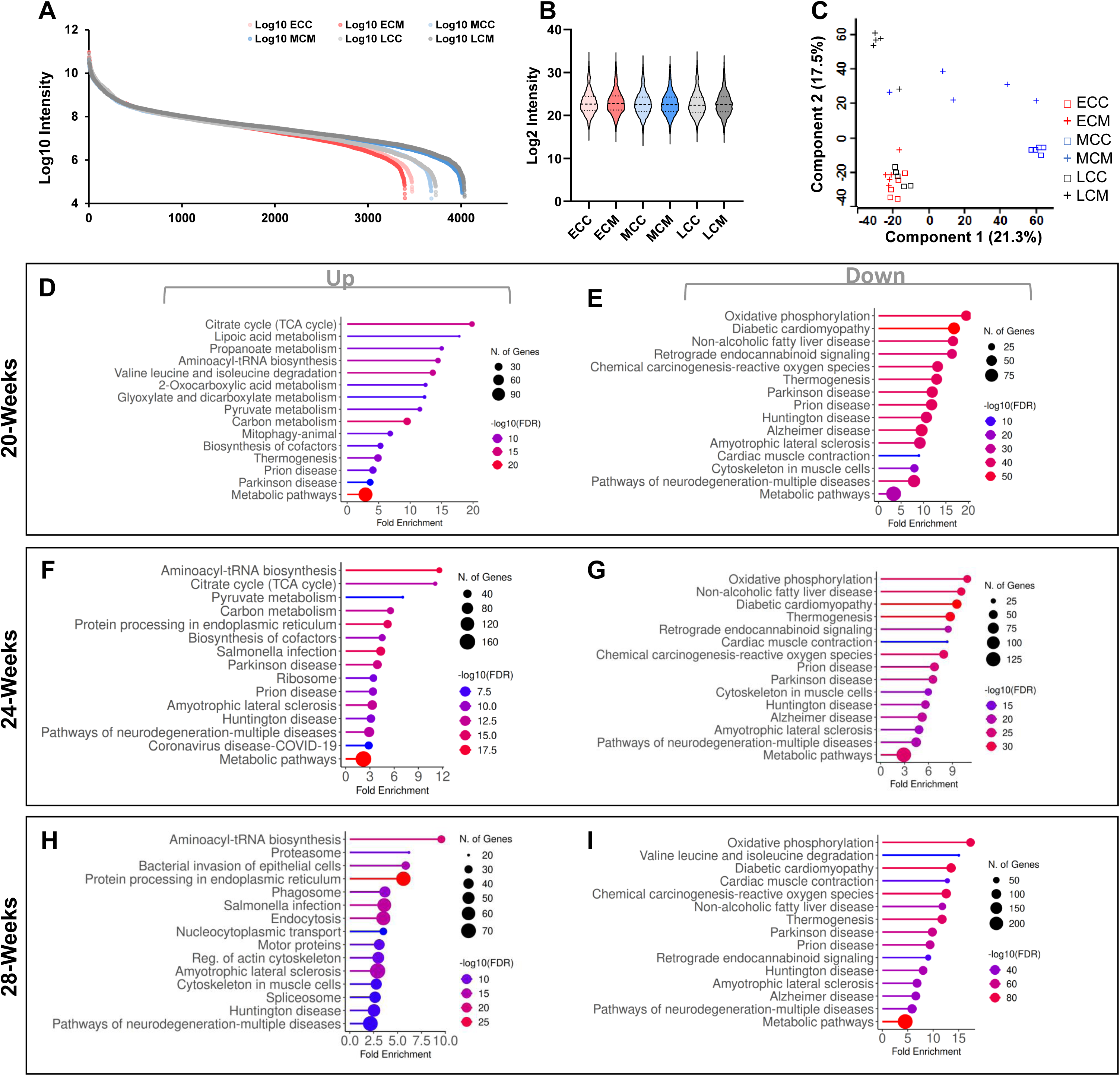
Damage to mtDNA in cardiomyocytes impacts proteomic pathways in the heart. **(A)** Log10 abundance rank of proteome detected within PolG^Mut^ and PolG^Con^ mice at 20, 24 and 28 weeks post-Tam and **(B)** Log2 violin plot of transformed intensity data showing data distribution (n=5/group/timepoint). **(C)** PCA reflecting the global proteome of each sample based on the log2 intensity transformed value of all samples. Gene ontology of the top 15 most significant KEGG pathway processes (FDR 0.15) upregulated at **(D)** 20 weeks **(F)** 24 weeks **(H)** 28 weeks post-Tam and downregulated at **(E)** 20 weeks **(G)** 24 weeks **(I)** 28 weeks post-Tam in PolG^Mut^ mice compared with PolG^Con^ mice. See methods for data processing and statistical analysis. Source data for these figures are provided in the Source Data file.

**Supplementary Figure 4:**
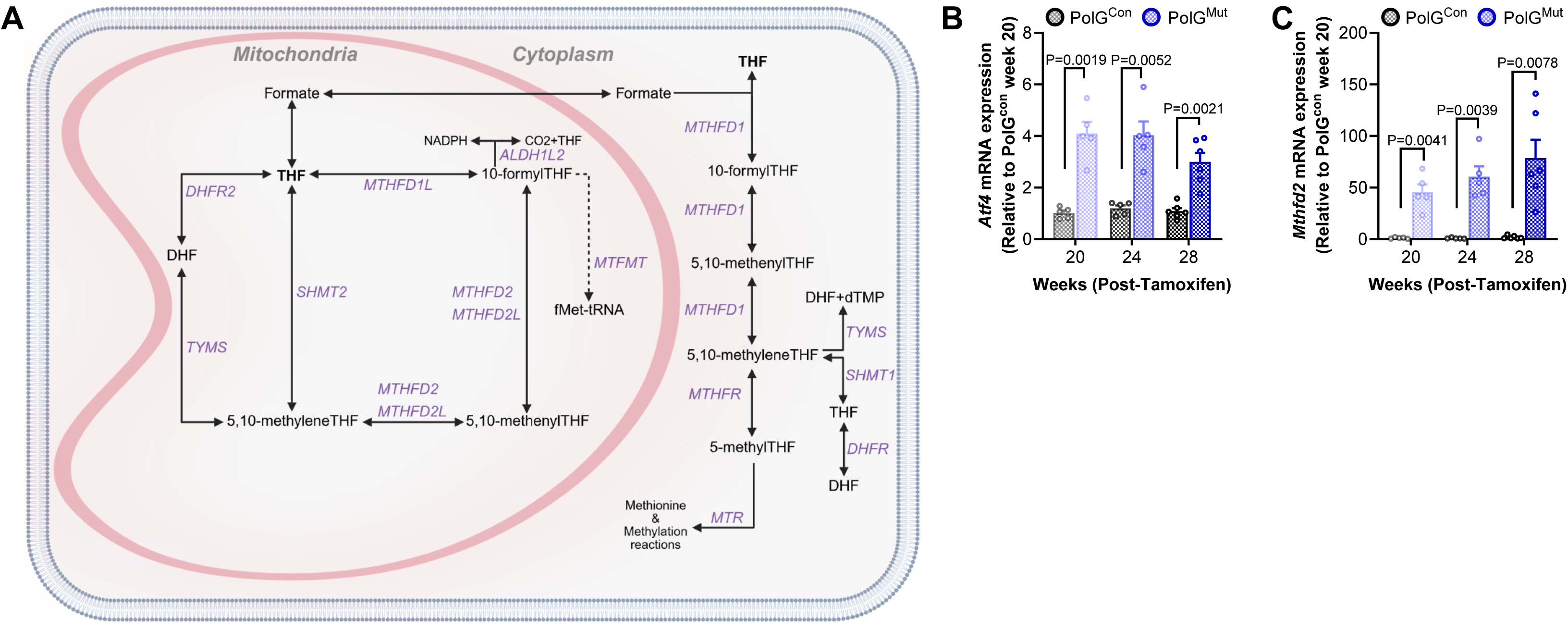
PolG mutation and subsequent mtDNA damage in cardiomyocytes activates the ISR and alters mitochondrial folate metabolism in the heart. **(A)** Schematic of folate metabolism compartmentalisation in the mitochondria and cytosol. **(B)** *Atf4* and **(C)** *Mthfd2* gene expression in LV from PolG^Mut^ and PolG^Con^ mice at 20 (n=5/group), 24 (n=5/group) and 28 (n=6/group) weeks post-Tam (relative to PolG^Con^ mice at 20 weeks post-Tam). Data are presented as mean ± SEM, with P-value between control and mutant biological replicates determined by unpaired welch t-test **(B, C)**. Source data for these figures are provided in the Source Data file. Fig S4A created in BioRender. Drew, **B.** (2025), http…….

**Supplementary Figure 5:**
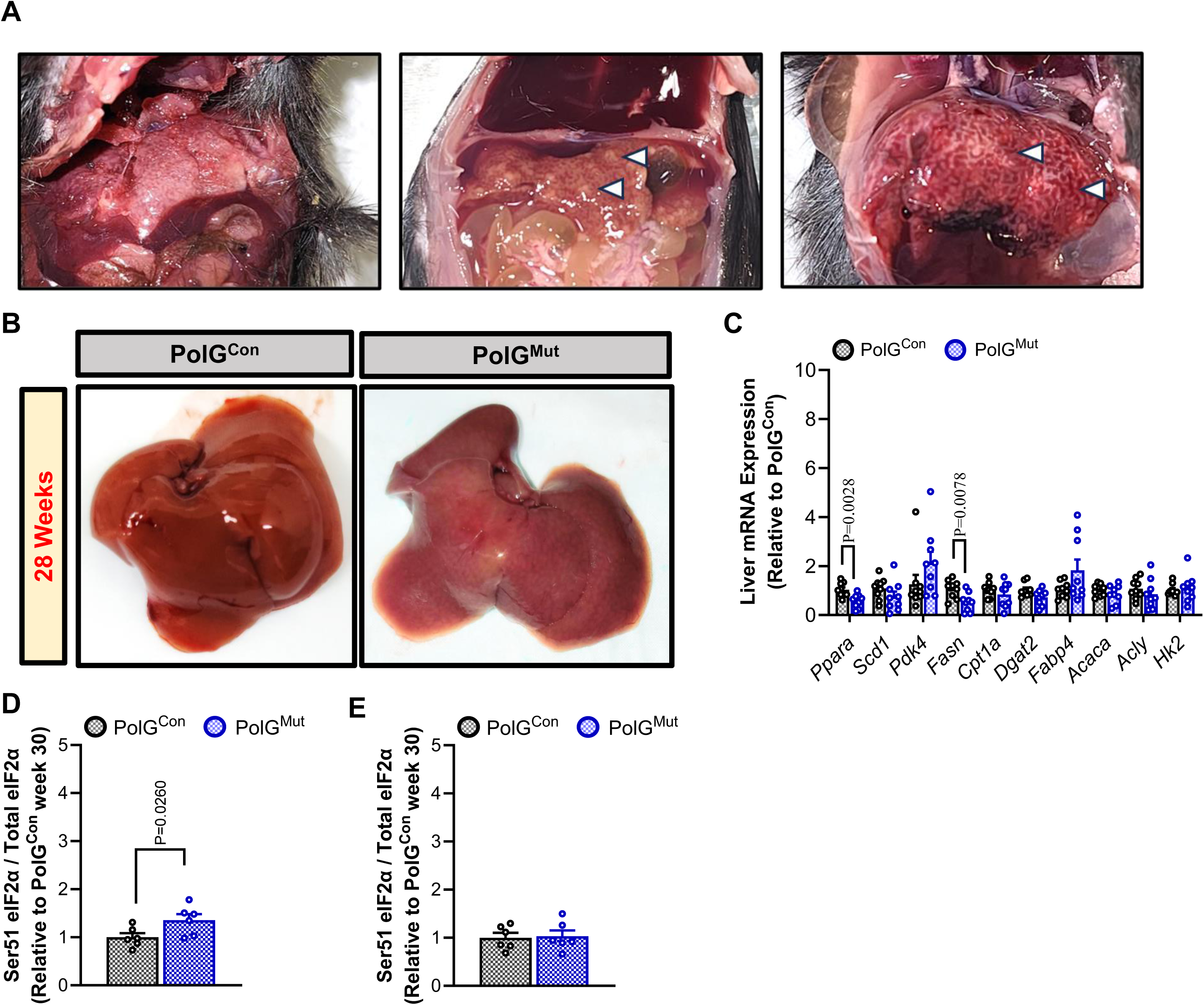
Cardiomyocyte specific activation of the ISR promotes cardiac hepatopathy. **(A)** Representative images of PolG^Mut^ mice livers at 30 weeks post-Tam, with white arrows indicating ‘nutmeg’ phenotype with cirrhotic nodules. **(B)** Representative comparative images between PolG^Mut^ and PolG^Con^ mice livers at 28 weeks post-Tam. **(C)** Metabolic gene expression (qPCR) in liver from PolG^Mut^ and PolG^Con^ mice at 28 weeks post-Tam (n=9/group). Data are presented as mean ± SEM, with P-value between control and mutant biological replicates determined by Mann-Whitney test. Quantitation of Western blots for p-eIF2α and eIF2α ratios in **(D)** adipose tissue and **(E)** Quadricep tissue. Source data for these figures are provided in the Source Data file.

## DATA AVAILABILITY

All raw and processed data is and will be made available in supplemental data and tables, and in online repositories where appropriate.

## ACKNOWLEDGMENTS

We acknowledge funding support from the Victorian State Government OIS program to Baker Heart & Diabetes Institute (BHDI). These studies were supported by funding from the BHDI Obesity & Lipid Program, as well as fellowship support to BGD from the National Heart Foundation of Australia (101789), to BGD from the NHMRC (Investigator Grant 2016530, Ideas Grant 2013158), and Grant support to STB from the National Heart Foundation of Australia (Vanguard Grant: 108166). We thank members of the MMA, Metabolomics and Molecular Proteomics laboratories at BHDI for their contributions. We are grateful for platform support from the Baker Preclinical Cardiology Platform, and the Monash Histology Platform at Department of Anatomy Monash University. Various figure panels were created using BioRender.com, with all such figure content sublicensed accordingly. Mitochondrial DNA sequence data was generated by CD Genomics (USA)

## AUTHOR CONTRIBUTION

BGD conceived the study and co-designed the experiments with STB. BGD and DCH conceived and generated the PolG floxed mouse model. BGD and STB wrote the manuscript. STB, YT, SW, SJ, CY, YL, KHL, HK and DGD performed animal experiments and phenotyping. STB, YT, SW, SJ, CY, YL, KHL, HK, DGD, JC, DWG, BGD analysed data, processed tissue samples, and performed molecular and biochemical experiments. DCH, DWG and BGD provided reagents, experimental advice, and access to infrastructure and resources. All authors read and approved the manuscript.

